# Evolutionary dynamics of the insertion sequence IS6110 in the *Mycobacterium tuberculosis* complex

**DOI:** 10.64898/2026.09.24.753865

**Authors:** Christoph Stritt, Chloé Loiseau, Sevda Kalkan, Sonia Borrell, Daniela Brites, Sebastien Gagneux

## Abstract

Insertion sequences (IS) are the most common type of transposable element in prokaryotes and shape the structure of genomes through transposition and by providing a substrate for recombination. Despite the mutational impact of IS, the evolutionary dynamics of most elements in most species remain unknown. Here we study the dynamics of IS6110 in 10,000 strains of the Mycobacterium tuberculosis complex (MTBC). We developed a tool that allows the detection and comparison of IS insertions from short reads without using a reference genome. Using ancestral state reconstruction (ASR) on presence-absence patterns of IS6110, we describe the distribution of copy numbers (CNs) in the MTBC, infer birth rates of the element, and identify genomic regions with large numbers of parallel IS6110 insertions. Copy numbers in the MTBC range from 1 in some clades to more than 30 in strains of La3 (M. orygis). IS6110 birth rates scale approximately linearly with copy number and are elevated on terminal branches, consistent with the delayed action of purifying selection. A key characteristic of IS6110 is its occurrence in hotspots: the 5% most frequently targeted regions account for half of all independent insertion events. The motif 5’-TCTCAAAW-3’ is enriched around target sites and in hotspots, suggesting that the accumulation of insertions in these regions results through a combination of non-random insertion and purifying selection in other regions. To conclude the study, we propose a niche constraints model according to which the distribution of IS6110 in the MTBC is governed by the rarity of regions that have both suitable DNA properties and little functional value for the host.

## Introduction

Transposable elements (TEs) are segments of DNA that can introduce copies of themselves in a genome through a variety of copy-and cut-and-paste mechanisms (Siguier et al., 2015; Wells and Feschotte, 2020). In most cases, they do not encode anything of direct use for the host. On the contrary, transposition tends to be disruptive (Consuegra et al., 2021; Darmon and Leach, 2014): it knocks out genes, disturbs regulatory networks, and through ectopic recombination induces deletions, inversions and duplications. All these mutation types can occasionally be beneficial (e.g. Hof et al., 2016; Vandecraen et al., 2017). In their ambiguous role as selfish replicators and sources of adaptive variation, TEs are thus an intriguing example of evolutionary conflict at the molecular level (Werren, 2011).

Insertion sequences (IS) are the most common type of TE in prokaryotes and consist of a transposase gene flanked by terminal inverted repeats (TIRs, Siguier et al., 2015). Prokaryote genomes may harbour from zero to few hundred IS copies, with genome size being the main correlate of IS abundance (Touchon and Rocha, 2007). Large genomes in prokarytes reflect increased coding capabilities that are required in complex environments (Kirchberger et al., 2020), and IS may find more available niches in large genomes without themselves contributing much to genome size (Touchon and Rocha, 2007). Despite the ubiquity and mutational impact of IS, little is known about most IS in most species beyond their DNA sequence. This reflects both technical limitations and conceptual preferences. IS are between 1,000 and 2,000 base pairs long (Siguier et al., 2015); if present in multiple near-identical copies, standard short reads are insufficient to assemble them. Mapping short reads against a reference genome, on the other hand, only yields “clean” insertion signatures in regions that are not repetitive and that are present in both the reference and the sequenced sample (e.g. Durrant et al., 2020; Hawkey et al., 2015). On the conceptual side, much attention has been paid to the functional consequences of transposition, whereas the evolutionary dynamics of IS are poorly known beyond broad generalisations from comparative genomics (Cerveau et al., 2011; Iranzo et al., 2014; Wu et al., 2015).

This study serves two purposes. Firstly, it introduces detettore6110, a tool that allows the detection and characterization of IS polymorphisms from short reads without using a reference genome. Secondly, the study investigates a specific IS from an evolutionary angle. We applied detettore6110 to 10,000 strains of the *Mycobacterium tuberculosis* complex (MTBC) in order to study the dynamics of IS6110—a comparatively well-known element that served as a sanity check for our tool and illustrates the gaps in our understanding of how IS evolve.

The bacteria of the MTBC are clonally evolving obligate pathogens that infect and cause tuberculosis disease in a range of mammalian species (Goig et al., 2025). In humans, tuberculosis remains a leading cause of death, and the ongoing evolution of multi-drug resistance is hampering efforts to control the disease (WHO, 2025). Variation in IS6110 copy numbers was first described in 1990 (Thierry et al., 1990) and subsequently used for strain genotyping in this low-diversity pathogen (reviewed by Merker et al., 2017). IS6110 is 1355 base pairs (bp) long, bounded by 28 bp imperfect terminal repeats and produces target site duplications (TSDs) of 3 to 4 bp upon integration. The sequence of IS6110 shows little variation within the MTBC (Thabet et al., 2015), and copy numbers range from zero (Lok et al., 2002) to more than 20 in lineage 2 (Shitikov et al., 2019) and La3 (also called *M. orygis*, Refaya et al., 2024).

Stress inducibility (Ghanekar et al., 1999; Safi et al., 2004), anecdotal evidence of beneficial insertions (Soto et al., 2004), and high IS6110 copy numbers in the globally successful lineage 2 (Shitikov et al., 2019) have fueled the view that IS6110 activity benefits the bacteria by increasing the evolvability of bacterial populations (Gonzalo-Asensio et al., 2018; McEvoy et al., 2007). This “evolvability hypothesis”, however, is typically framed in an adaptationist spirit that fails to consider alternative evolutionary scenarios (Koonin, 2016). Actual evidence that IS6110 benefits the bacteria is scarce, and one study that explicitly compares different evolutionary models suggests that purifying selection largely explains IS6110 copy number distributions (Tanaka, 2004).

Here we place patterns of IS6110 variation in a phylogenetic context and focus on the evolutionary dynamics of IS6110. More specifically, we 1) describe the distribution of copy numbers in the MTBC, 2) reconstruct ancestral states and model IS6110 birth rates along the phylogeny, and 3) infer the number of parallel insertions in hotspot regions and target site preferences. Uniting these disparate aspects, we propose a niche constraints model according to which observed patterns such as copy number variation and insertion hotspots reflect the rarity of genomic regions that both offer suitable target-site properties and impose little fitness cost on the host.

## Results and Discussion

### Reference-free detection of IS polymorphisms

To avoid reference bias and make better use of the information in short reads, we developed a method that allows to infer IS polymorphisms and copy numbers independently of a reference genome. The method, implemented in detettore6110, is outlined in figure 1 and described in detail in the methods section. The basic idea is that insertion sites are distinguished from each other not by their position relative to a reference genome, but by the DNA sequences flanking the insertion (Fig. 1A). Flanking sequences (FS) are identified by mapping reads against an IS family consensus sequence. Subsequent read clustering and local assembly produce 5’ and 3’ consensus flanking sequences (cFSs) that provide, given some limitations detailed below, a precise identifier for an insertion site. By comparing the cFSs of different strains, unique 5’ and 3’ flanking sequences (uFSs) are identified and presence-absence matrices are constructed (Fig 1B). Combined with a phylogenetic tree, these provide the basis to study the evolutionary dynamics of the IS.

**Figure 1:**
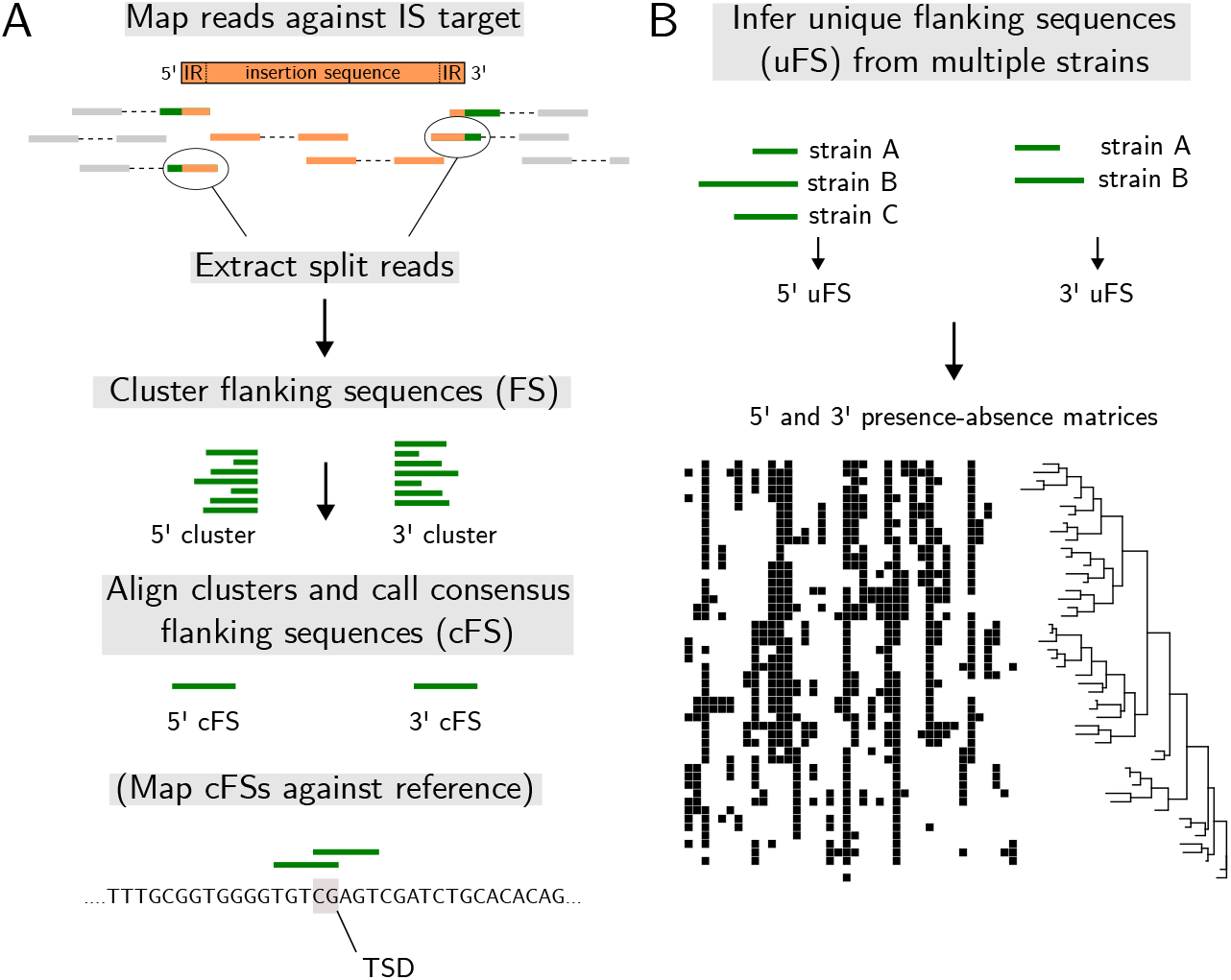
Workflow of detettore6110. A) Identification of insertion sites in a single strain, with the optional step of mapping consensus flanking sequences (cFS) against a reference genome. In the schematic depiction of the insertion sequence, IR denotes the inverted repeats that mark the beginning and end of the IS. B) With results available for multiple samples, the cFS of different strains are combined into a matrix of unique flanking sequences (uFS).

To benchmark our method, paired-end reads were simulated at different lengths and coverages from 28 diverse MTBC assemblies (Supplemental Table S1). IS6110 was annotated in the assemblies and the regions flanking the insertions were extracted and compared to the cFS produced by detettore6110. In total, 326 IS6110 copies in 28 assemblies were used for benchmarking. The benchmarking results show sensitivity surpassing 99% at coverages of 40-fold and higher (Fig. S1).

The only combination that performed markedly worse (<90%) was 20-fold coverage with 100 bp reads, whereas 150 bp reads with 20-fold coverage performed well (> 99%). Precision offers a different picture (Fig. S1): false positives were produced by some assemblies independently of read length or coverage. A closer inspection of what caused them showed that the algorithm had detected IS6110-specific inverted repeats (IRs) that were not part of a full element. In total only six such IRs were present, and they caused low precision in strains with low copy numbers: the lowest precision (0.5) resulted from a strain with two full IS6110 copies and two false positives due to solitary IRs.

In summary, our method allows to identify IS polymorphisms at high sensitivity and precision. A principal limitation is that IS presence is inferred from reads reaching into the element from either side but not spanning the full element. This means that solitary IRs cannot be distinguished from intact elements. It also means that the 5’ and 3’ flanking sequences of a given insertion can not be linked, such that for a single insertion there is independent evidence for the 5’ and 3’ junctions.

### Large copy-number variation between and within lineages

We next applied our method to 10,000 published genomes that represent the known diversity of the MTBC (Table 1, Supplemental Table S2). Strains were selected based on the quality of the sequencing data, and well-represented lineages were downsampled while we kept as many strains as possible of understudied lineages, in particular those associated with animals (see Methods). Based on the sequences flanking the IS6110 insertions, we constructed presence-absence matrices showing which insertion sites are shared between which strains. 10,102 uFSs based on the 5’ and 10,071 based on the 3’ cFS were identified. A main characteristic of these matrices is that they are sparse: the majority of the insertion sites are occupied once in a single strain (5’: 6026, 3’: 5945), and the site frequency distribution shows a strong skew towards rare alleles (Fig. S2).

**Table 1.**
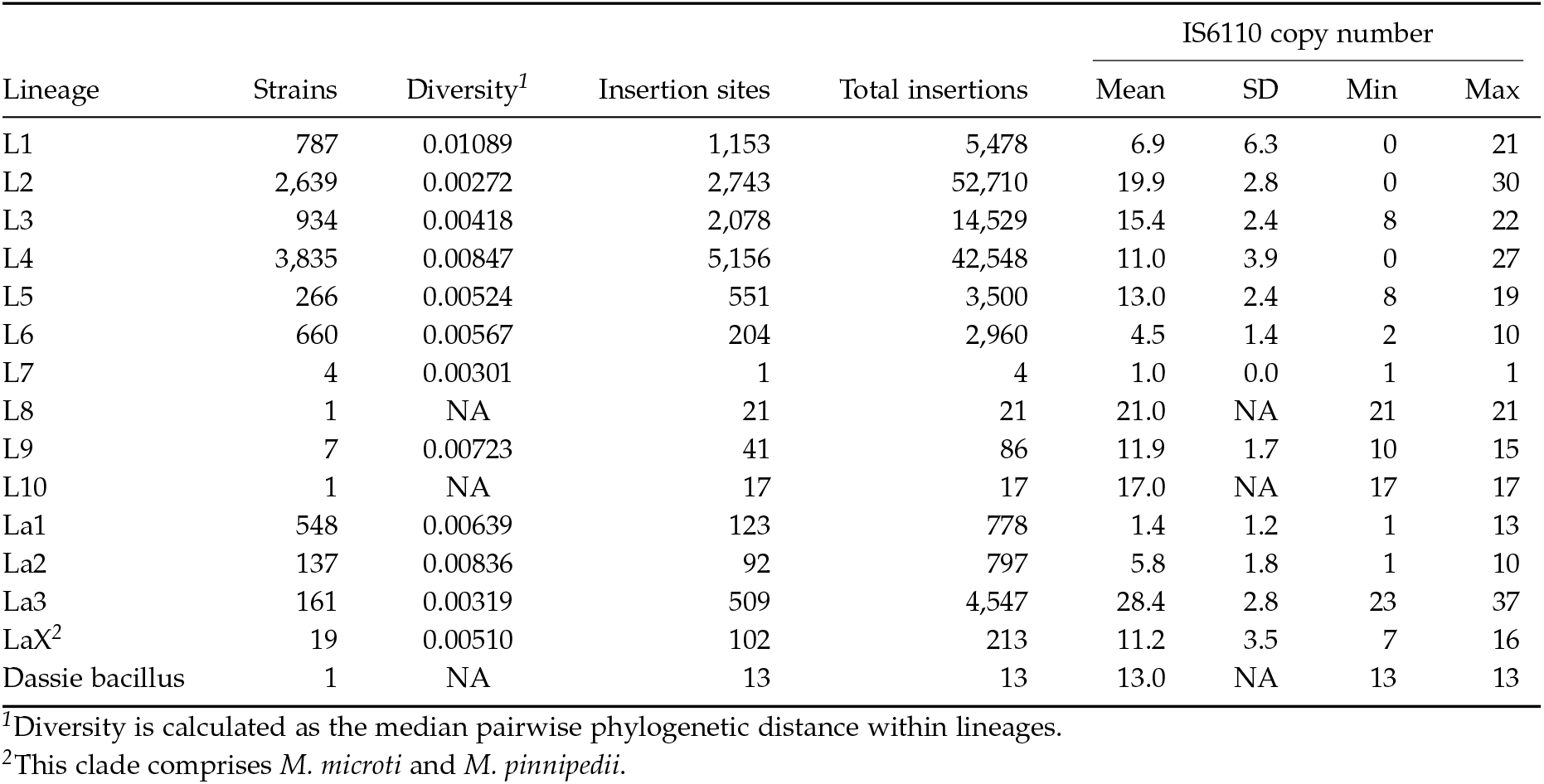
IS6110 insertions in 10,000 strains.

To provide a first overview of IS6110 variation in the MTBC, and as a sanity check for our IS detection approach, we estimated IS6110 copy numbers (CN) conservatively as the smaller of the 5’ and the 3’ cFS counts per strain. The 5’ and 3’ cFS counts correlate strongly (*r* = 0.995, Fig. S3), suggesting that solitary IRs are not an issue in real data and that IS6110 is mostly intact. In addition, we inferred a phylogenetic tree from single nucleotide polymorphisms and used it to reconstruct ancestral CNs through maximum parsimony, which allowed us to assign a CN to each internal node of the tree. Figure 2 provides a global picture of IS6110 copy number distribution in the MTBC. CNs range from one in L1, La1 and the four L7 strains to a mean of 20 in L2 and of 28 in La3 (Fig. 2, Table 1). Only 16 strains were found to contain no IS6110 insertion, 14 of them belonging to L1 and 9 forming a single clade in L1 (L1.1.1.1). While the L4 strain with 0 copies belongs to a low-CN clade within this lineage (L4.2.2, mean CN = 2), the zero-copy L2 strain is difficult to reconcile with its phylogenetic position. Re-inspecting the reads from this strain, we observed that the number of reads mapping to internal parts of IS6110 is comparable to other L2 strains from the same Bioproject; at the ends, however, read coverage drops sharply, resulting in very few reads that span the insertion breakpoints (Fig. S4). As this pattern is an outlier (Fig. S4), we suspect that it reflects a sample-specific technical artefact, possibly caused by read trimming or preprocessing.

**Figure 2:**
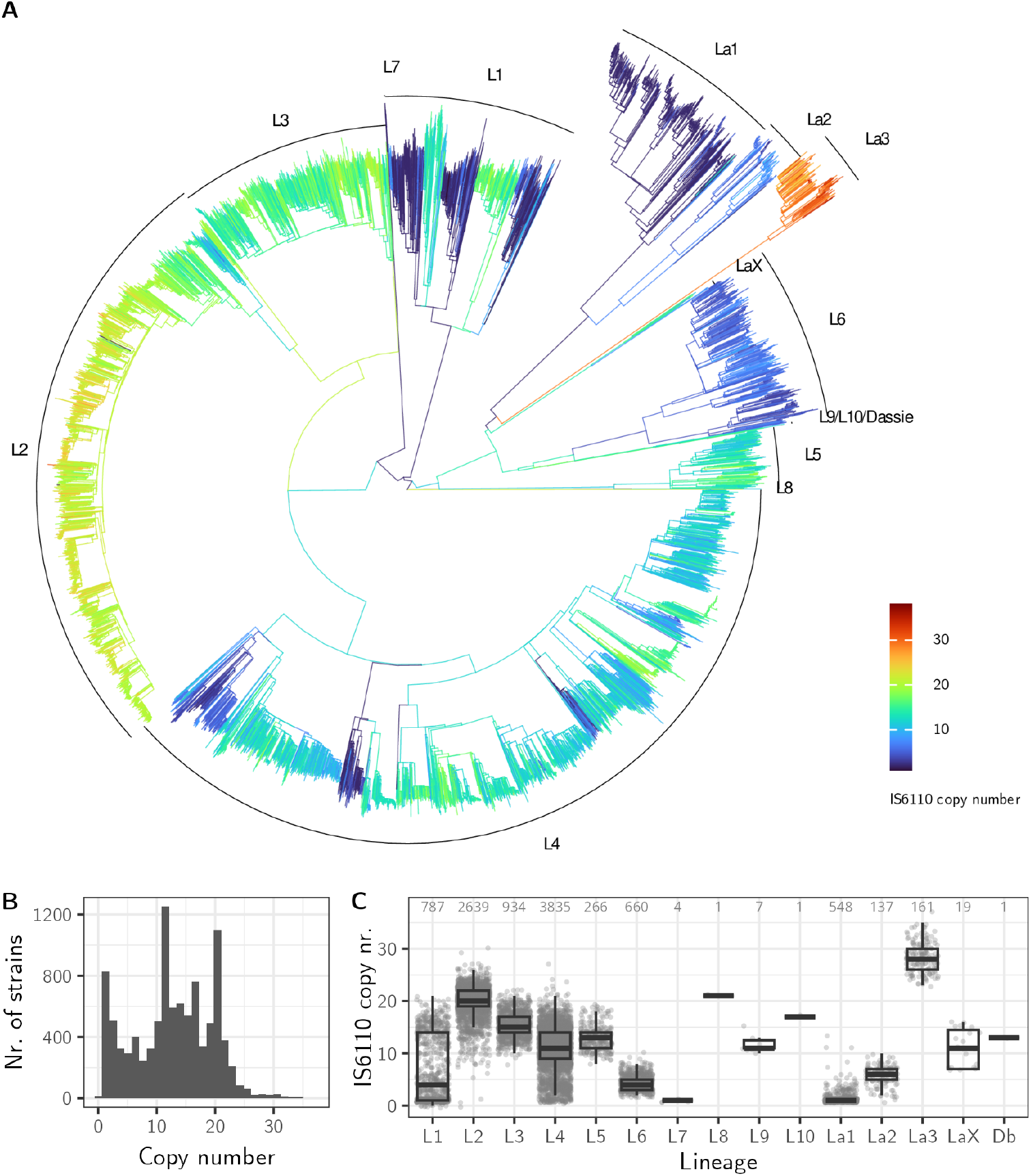
Copy number variation in the Mycobacterium tuberculosis complex. A) Rooted maximum-likelihood phylogeny of the 10,000 strains, with tip colors representing IS6110 copy numbers. B) The multi-modal distribution of IS6110 copy numbers in the MTBC. C) Distribution of IS6110 copy numbers in the different lineages. The number of strains included for each lineage is shown on top.

CN distribution shows a strong phylogenetic signal that does not necessarily correspond to lineage boundaries. The standard deviation of copy numbers is highest in lineage 1 (Table 1), where two sublineages (L.1.1.3 and 1.2.1) have evolved intermediate CNs whereas strains belonging to L1.1.1, L1.1.2 and L1.2.2 mostly have one or two copies. The second highest variation in CN is observed in L4, which covers much of the spectrum from low (<5) to high (>20) CN and where stark contrasts occur between sister clades. A considerable range of CNs was found in all lineages that had more than a few samples (Fig. 2A), testifying to dynamic IS6110 landscapes across the MTBC.

The observation of IS6110 CN variation is not new: since the early 1990ies, IS6110-based restriction fragment length polymorphisms (RFLPs) have been used to genotype MTBC strains (Thierry et al., 1990) and a large, fragmented literature describes the occurrence of IS6110. Furthermore, recent studies based on whole genomes have investigated differences between human-associated lineages (Gonzalo-Asensio et al., 2018; Roychowdhury et al., 2015), with a particular focus on L2 and the potential functional impact of insertions (Alonso et al., 2013; Antoine et al., 2021; Shitikov et al., 2019). This first part of our study is thus largely confirmatory and shows that the approach implemented in detettore6110 is sound. Beyond the confirmatory, a global phylogenetic view on IS6110 copy number variation (Fig. 2) highlights patterns that have received scant notice. Firstly, La3 strains, which infect a range of animals including cattle, camels, buffalos, deer and humans (Hugh et al., 2025), have the highest CNs and define a new upper boundary at 37 copies. A recent study has hinted at this scenario (Refaya et al. 2024), which disrupts the mantra of L2 as the lineage with most copies and may provide new insights into the implications of high CN. A second intriguing pattern is the extensive CN variation in lineage 4. Previous studies have emphasized differences between lineages (Gonzalo-Asensio et al. 2018), but variation within lineages is generally present and sometimes large. CN variation has some practical relevance. IS6110 is MTBC-specific and its amplification underlies different diagnostic tests (e.g. Kechin et al., 2023). Strains with zero copies are rare (Fig. 2B) and most of them belong to sublineage L1.1.1.1. More common are strains with a single copy, as shown by the first peak in multimodal CN distribution (Fig. 2B). Some low-CN clades are prevalent in some regions, and in these contexts the sensitivity of IS6110-based diagnostics may be lower than expected. This particularly applies to sublineages of L1: L1.1.1 strains are prevalent in countries of mainland Southeast Asia (Cambodia, Laos, Vietnam, Thailand), L1.1.2 strains in India, Pakistan and countries in East Africa, while L1.2.2 strains cause many infections in South Asia as well as East/South Africa (reviewed by Rojas et al., 2025).

### Pairwise IS6110 differences reflect phylogenetic depth and copy numbers

Copy numbers themselves reveal little about the diversity caused by IS6110. Between two strains with ten copies, for instance, how many copies are conserved and reflect the same insertion events in a common ancestor? And how many reflect different insertions or loss of a previously shared copy? To make better use of the IS6110 presence-absence patterns, we first counted by how many IS6110 uFSs strains differ. Figure 3 shows the distribution of pairwise differences *d*_IS_ within lineages. Ranges are generally large, given that lineages include near-identical to more distantly related strains. At the extremes, strains within L4, L2, and La3 may differ by up to 40 copies, while strains within the low-CN lineages L6, L7, La1 and La2 differ by few to no copies. IS6110 diversity is highest in L4 (median *d*_IS_ = 21), followed by the composite lineage LaX (*M. microti, M. pinnipedii*) with its deep branches, L5 and L3.

**Figure 3:**
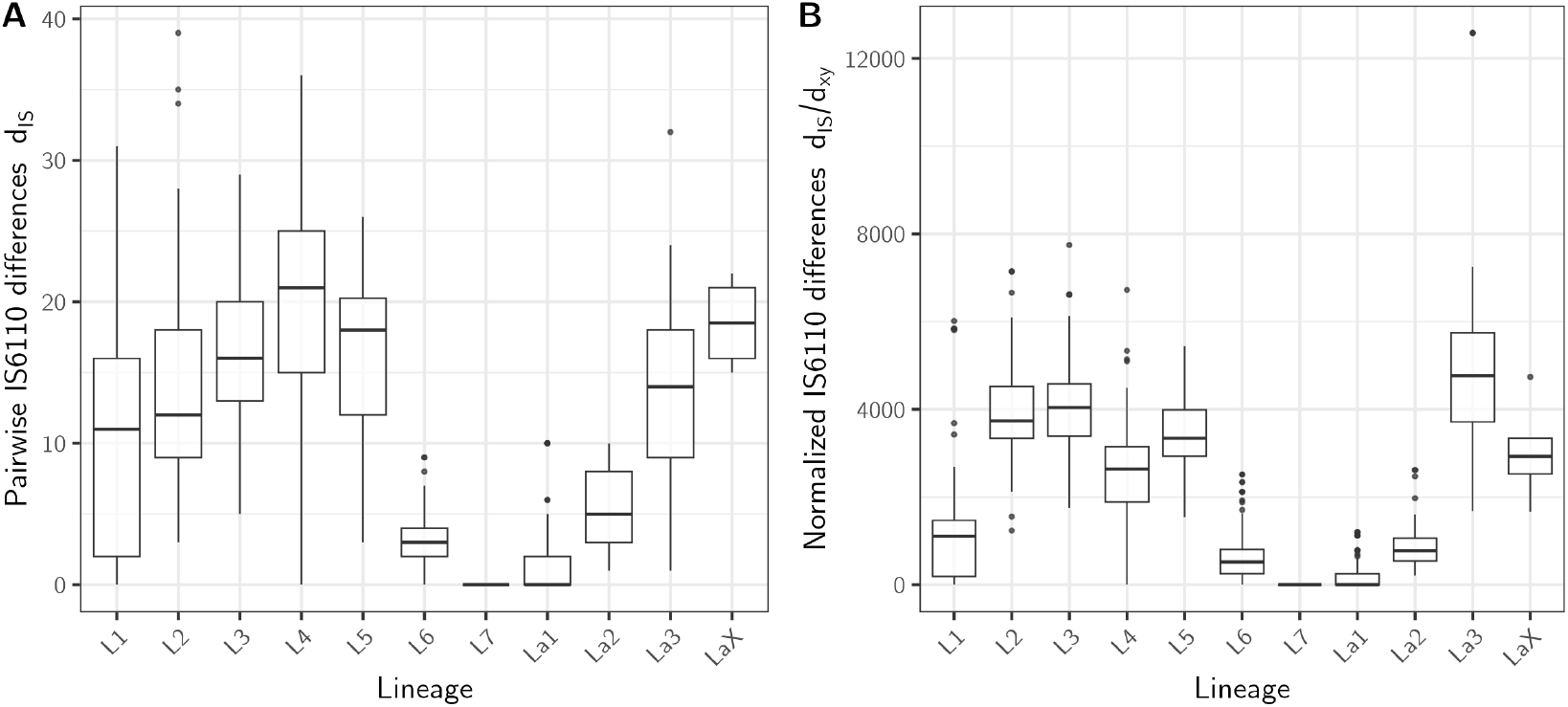
A) Distribution of the number of pairwise IS6110 differences between strains within lineages. B) Distribution of normalized IS6110 differences between strains. Normalization was achieved by dividing d_IS_ by the pairwise phylogenetic distance d_xy_, which was extracted from the tree in figure 2. To decrease the computational burden of pairwise comparisons, well-represented lineages were downsampled to a maximum of 1500 strains. Lineage LaX includes M. microti and M. pinnipedii.

High CNs do not translate into particularly high *d*_IS_: L2, the human-associated lineage with the highest mean CN, is less diverse than other lineages. This is because a comparatively large proportion of IS6110 insertions are present at high frequency in L2 (Fig. S5): 15 insertion sites are shared among 75 to 100% of all L2 strains. In La3 this pattern is even more extreme: 24 sites are in the 75-100% frequency category. In contrast to these high-CN lineages, not a single of the 5156 sites in L4 is present in more than 75% of the strains, and only a single site is shared among 50 to 75% of the strains (Fig. S5). L4 is genetically more diverse than L2 and La3, has deeper structure and less shared history (Fig 2A, Table 1). Not too surprisingly, these phylogenetic differences are also reflected in the frequency spectrum of IS6110 insertions. When normalizing *d*_IS_ by pairwise phylogenetic distance *d*_xy_, high-CN lineages do have the highest IS diversity per unit base substitution (Fig. 3B).

In summary, genetic variation caused by IS6110 can be large within and, by extension, between lineages. In lineages with intermediate to high CN, it is common to find strains that differ by 15 to 20 insertions (Fig. 3A). Lineages with high CN are not particularly diverse in absolute terms because they happen to be phylogenetically shallow. But they do have the most dynamic genomes per unit substitution/time. Put differently, high IS6110 copy numbers increase the relative mutational burden of IS compared to single nucleotide substitutions.

### Copy number and purifying selection affect IS6110 birth rates

Because IS vary in copy number, their contribution to genetic diversity is inherently more heterogeneous than that of point mutations. To better understand the extensive variation of IS6110 activity in the MTBC, we used ancestral state reconstruction (ASR) to infer the birth and death of copies along the branches of the phylogeny, and linear mixed effect models in order to disentangle factors contributing to variation in birth rates. Ancestral states of IS6110 presence and absence were inferred for each uFS using maximum parsimony, as for the copy numbers above, and the number of 0*→* 1 and 1 *→* 0 transitions were counted on each branch.

In total we inferred 17,579/17,289 births and 4008/3584 deaths of IS6110 along the tree from the 5’ and the 3’ uFS matrices, respectively. Independent estimates from the 5’ and the 3’ uFS show a high correlation for the number of births on each branch (*r* = 0.95, Fig. S6), whereas the correlation for deaths was substantially lower (*r* = 0.65, Fig. S6). Maximum-parsimony ancestral state reconstruction has known limitations in inferring loss events, as uncertainty in observed character states is not modeled and translates asymmetrically into an excess of inferred losses relative to gains (Cunningham, 1999). In the following, we focus on the dynamics of IS6110 births and outline additional approaches that could be used to study IS6110 deletion dynamics. Analyses were conducted with both the 5’ and the 3’ matrices; since the outcomes agreed, for simplicity we here only report the results obtained from the 5’ matrix.

In agreement with the prevalence of singleton insertions (Fig. S2), 75% of the births occurred along the terminal branches of the tree. Most branches exhibited few or zero births and a small subset show many births (Fig. S7A). The number of births scales with branch length, though in a non-trivial way with many exceptions (Fig. S7B). This is expected since birth rates depend on IS6110 copy numbers (Fig. S7C), and few insertions may occur on a long branch if there are few copies present. To better understand the interplay of these factors, we modelled the number of births on a branch as a function of copy number at the parental node and branch type (terminal vs. internal). A generalized linear mixed model was fitted where we allowed independent intercepts and slopes for each lineage in order to account for the strong lineage-structure in the distribution of copy numbers (Fig. 2). Branch length was used as an offset in the model and we accounted for the non-independence of branches that share a parent node by including parent node as a random effect (see Methods). Parental copy number was log-transformed such that a potential non-linear effect on birth rates, reflecting self-inhibition or positive feedback, could be captured.

Our model suggests that IS6110 birth rate scales approximately linearly with copy number, with an estimated exponent *β* = 1.06 (± 0.13 SE) close to unity (Fig. 4). The position of the branch in the tree also affected IS6110 birth rates: terminal branches exhibited higher IS6110 birth rates than internal branches (log rate ratio = 0.135 ± 0.021 SE), corresponding to an approximately 1.15-fold increase in expected birth rate on terminal branches (Fig. 4). An example: the expected IS6110 birth rate in an ancestral strain with 10 copies is 0.00055 insertions per base substitution per genome on terminal branches and 0.00048 on internal branches; an additional copy increases these rates to 0.00061 and 0.00053, respectively. This corresponds to median predicted per-copy rates of 4.89 *×*10−^5^ births per substitution per genome on internal and 5.60 *×*10^*−*5^ births on terminal branches. Lineage-level random intercepts (SD = 0.74) and slopes varied (SD = 0.26) and were negatively correlated (r = *−* 0.97), meaning that lineages with higher baseline birth rates tended to exhibit weaker copy-number dependence (Fig. S8).

**Figure 4:**
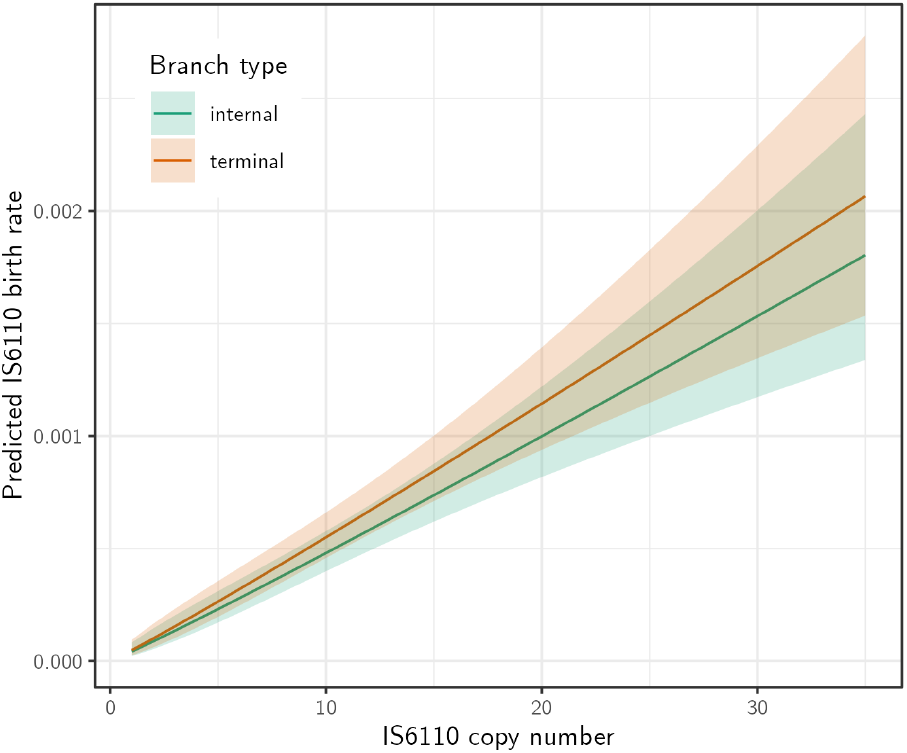
Effects of IS copy number and branch type (terminal/internal) on IS6110 birth rates. Lines and confidence intervals show the lineage average effect of copy number separately for internal and terminal branches. The rate is in births per base substitution per genome: 0.001, for instance, means one expected IS birth per 1000 base substitutions.

Whereas the transposition rate describes the rate at which elements actually insert (analogous to the mutation rate), the IS6110 birth rate is the slower rate at which elements were maintained during the evolution of the MTBC after having been affected by natural selection and genetic drift (analogous to the substitution rate, see Ho et al., 2015). IS6110 transposition rates have previously been estimated from serial isolates: Tanaka, 2004 reported a per-element transposition rate of 7.9 *×* 10^*−*5^ per site per generation. Assuming a range of clock rates (10^*−*6^, 10^*−*7^, 10^*−*8^) and generation times (18, 24, 30 hours) that cover values reported in the literature (Menardo et al., 2019; Stritt and Gagneux, 2023), our rate translates into per-element rates between 2.65 *×* 10^*−*7^ (slow clock rate, slow generation time) and 3.54 *×* 10^*−*9^ (fast clock, fast generation time) per site per generation. These estimates are two to four orders of magnitude smaller than Tanaka’s transposition rate, indicating that strong purifying selection filters most new IS6110 insertions before they reach observable frequencies. Most transposition events are either lethal or deleterious enough to prevent their carriers from founding successful lineages.

We have not come far in understanding the second key event in the life history of IS6110, its “death” through inter-element recombination. More sophisticated ancestral state reconstruction approaches that model error rates explictly could be tried on smaller data sets in order to infer death events more reliably. In addition, further mechanistic insights into deletion could be obtained by scrutinizing the short reads for deletion signatures such as the absence of TSDs, which is expected after homologous recombination between elements.

### Extensive parallel disruption through IS6110 insertions

An intriguing characteristic of IS is their occurrence in hotspots. IS6110 hotspots have been noted early on in genotyping studies (Fang and Forbes, 1997; Kurepina et al., 1998). In the absence of a phylogeny, however, shared ancestry has not been distinguished from independent insertion events as an explanation why some regions frequently carry IS6110. The extent of parallelism thus remains unknown. In this section we use ASR to infer how frequently independent disruptions of the same regions have occurred. Our reference-free IS detection approach does not provide genomic coordinates, but these can be obtained by mapping the cFS against a genome assembly and extracting insertion signatures in the form of overlapping 5’ and 3’ cFS (Fig. 1A, see Methods). As reference we used the reconstructed ancestral reference genome MTBC0, which contains several large regions that were lost in some lineages and is less affected by reference bias (Harrison et al., 2024). 8564 unique reference insertion sites were identified with expected target site duplications of three or four bp. This corresponds to 85% of the insertions inferred without using a reference.

IS6110 is clearly not distributed in a random way (Fig. S9). Copies of IS6110 are 4.14-fold overrepresented in intergenic regions (Fisher’s exact test, 95% CI 3.96-4.33, Fig. S9B). This is more pronounced when considering shared insertions (95% CI 3.97-4.58) rather than insertions occurring in single strains (95% CI 3.82-4.29), consistent with stronger purifying selection in genic regions and delayed selection against recent insertions. Of the 4012 genes annotated in the MTBC0 genome, 873 were at least once disrupted by an IS6110 insertion, while 459 out of 3166 intergenic regions were affected. Only four genes that were described as essential *in vitro* (DeJesus et al., 2017) were interrupted by IS6110: *Rv2017*, a transcriptional regulator that was knocked out in seven strains belonging to L5 and L2.2.1; as well as *Rv0013, Rv2357c* and *Rv3222c*, which were all disrupted once in a single strain. On the other hand, 2328 genes that appear to be non-essential for growth *in vitro* were devoid of IS6110 copies. This is a maximum estimate: on the one hand, 15% of the insertion sites could not be mapped to the reference. On the other hand, an accumulation analysis shows that our “natural mutagenesis experiment” was not saturated: adding more strains would result in additional disrupted genes, even in well sampled lineages (Fig. S10).

More informative than the mere presence or absence of insertions in genic and intergenic regions is the frequency of disruptions. We again resorted to ASR in order to estimate the number of times a region was disrupted independently. All insertions with mappable reference positions were used and aggregated per region, such that for each genic and intergenic region the number of IS6110 births could be inferred. The results show the characteristic skew towards rare events (Fig. 3A): of the 1332 regions affected by IS6610, 505 were disrupted only once, 386 thereof in a single strain. 1082 regions showed fewer than 10 parallel disruptions, whereas 154 regions had more than 20 and 62 regions more than 50 independent insertion events. To put a number on how hotspot regions contribute to and constrain IS6110 dynamics, we calculated the Gini coefficient, a measure of distributional inequality typically applied to income and wealth. When applied to the number of independent births contributed by each of the 1332 regions with IS6110, we inferred a Gini of 0.75. This is higher than the income inequality in any country: the 5% most frequently targeted regions account for half of all independent birth events, while the remaining 95% of regions collectively account for the other half (Fig. 5B). The top five percent includes 68 regions with at least 48 independent IS6110 births in each, which in turn corresponds to ca. 1% of all genic and intergenic regions of the genome.

**Figure 5:**
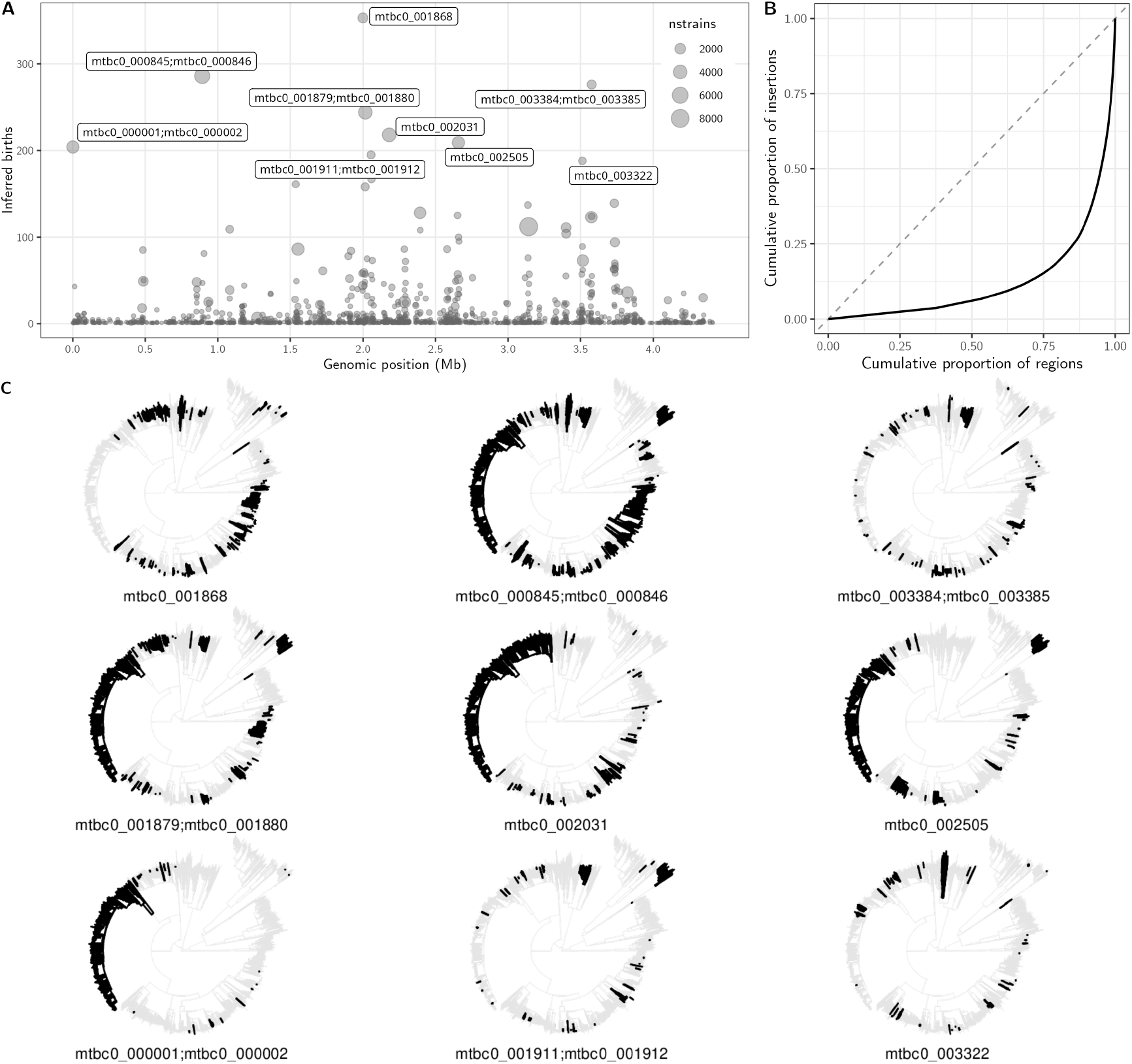
A) Genomic distribution of IS6110 hotspots, showing on the y-axis the number of parallel insertion events inferred through ancestral state reconstruction. Point sizes indicate the number of strains in which the region is disrupted. The nine top hotspots are labelled. Orthologous gene names in the reference strain H37Rv are provided in tabular summary of insertion hotspots in Supplementary Table S3. B) Lorenz curve, showing the cumulative contribution of IS6110-carrying regions to the total proportion of insertions. The dashed diagonal shows the scenario where each region contribites equally to the total amount. The Gini coefficient is the area between the dashed and the solid line (0.75); it would be 0 under full equality and 1 under maximum inequality (one region contributing all insertions). C) The nine hotspot regions with the highest numbers of independent insertions. The phylogenies are the same as in Fig. 2, dark branches show the strains and clades in which a particular region is disrupted.

At the high end of the spectrum are the insertion hotspots (Table 2, Fig. 5A,C). The top hotspot, with 353 inferred independent insertions and disruption in 1580 strains, is the phospholipase C gene *plcD* (*mtbc0_1868*). Second is the intergenic region *mtbc0_000845;mtbc0_000846* that, on closer inspection, overlaps with a copy of the insertion sequence IS1547 (Fig. S11). RFLP studies already noted this region as hypervariable and accordingly named it “IS6110 preferential locus” (ipl, Fang and Forbes, 1997). Among the top hotspots are furthermore different PPE genes, the *dnaA;dnaN* region that contains the origin of replication, transcriptional regulators (*mtbc0_001459, mtbc0_003384*), and genes of unknown function. As illustrated in figure 5C, the extent of parallelism in these regions is remarkable and reflects the lineage differences described above: five of the top nine regions are more or less consistently disrupted in L2 and four in La3, while L4 is characterized by multiple independent disruptions.

**Table 2.**
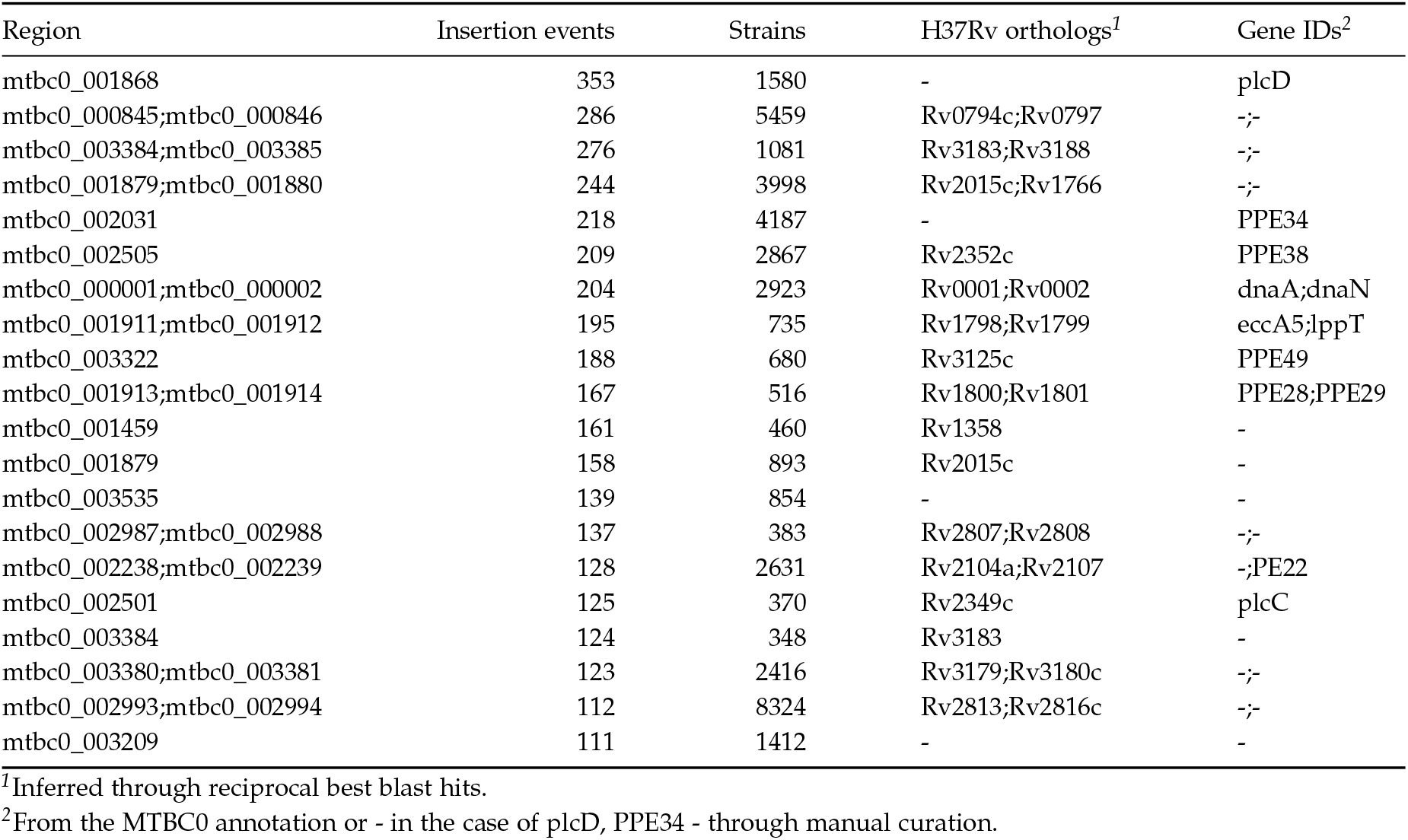
IS6110 insertion hotspots.

Other interesting regions were not among the top hotspots, but still showed some degree of parallelism. Upregulation of *phoP* through upstream insertions of IS6110 is the most frequently cited example of IS6110-mediated adaptation: changes in virulence traits regulated by this pleiotropic gene might have allowed usually zoonotic La1 strains to spread in a human population (Gonzalo-Asensio et al., 2014; Soto et al., 2004). In our data set, we identified 40 independent insertions in the intergenic region upstream of *phoP*. The majority of the 307 affected strains, 257, belonged to the L2.2.1.1/Pacific_RD150 clade, 21 to lineage 3, and the rest were spread across single or few strains in L1, L4, L5, L6, La1, La2. Genes involved in antibiotic resistance, which provide the most striking examples of convergent evolution in the MTBC, showed limited parallelism (Table S3). Using the WHO catalogue of resistance mutations as a reference, we overall identified 42 strains with insertions into genes associated with resistance. There were eight independent insertions into *mtbc0_002101* (*Rv1979*) and six into *mtbc0_000717* (*Rv0678*)—genes linked to resistance against bedaquiline and clofazimine (Zhang et al., 2015)—as well as a single disruption of *pncA* (pyrazinamide resistance) and two disruptions of *ethA* (ethionamide). The best documented example, *Rv0678*, is a repressor of the MmpS5–MmpL5 efflux pump, whose disruption causes increased drug export (Sonnenkalb et al., 2023). We conclude that, while biologically plausible, IS6110 insertions causing loss of function are rare compared to indels and missense mutations in resistance genes.

As this analysis was based on reference positions, we have certainly missed hotspots in more dynamic regions of the genome. But they are unlikely to affect the overall picture: hotspot regions are a pervasive characateristic of the IS6110 landscape, and some genomic regions have been disrupted dozens of times independently by IS6110.

### IS6110 prefers short, AT-rich motifs

Hotspot formation may reflect three non-exclusive processes: target site preference of the IS, convergent positive selection on insertions in certain genomic contexts, and relaxed purifying selection permitting accumulation in functionally tolerant regions. To our knowledge, while some suggestive vocabulary has been used to refer to IS6110 hotspots (e.g. “preferential insertion regions” in Roychowdhury et al., 2015), no study has so far investigated target site preferences of IS6110. To fill this gap, we joined the 8564 5’ and 3’ flanking sequences with mappable reference positions and used the STREME motif discovery algorithm to identify motifs enriched at insertion sites. Several motifs were enriched relative to genome-wide random windows of matching sizes (Table S4). The top motif, 5’-TCTCAAAW-3’, was detected in 40.3% of the flanking sequences (versus 11.5% of the background sequences) and showed a symmetric enrichment around the insertion site, with peaks 22 base pairs upstream and downstream of the center (Fig. 6). Repeating the analysis with 10 different sets of random background sequences showed similar top motifs with some small variation in degeneracy and motif length and the same symmetry around the insertion site (Fig. S12). The positional symmetry suggests that the transposase prefers an AT-rich sequence context centered approximately 22 bp upstream and downstream of the insertion site. This could reflect a preference for physical properties of the DNA such as bendability or minor groove width, which correlate with AT-rich sequences (Chandler and Mahillon, 2007; Craig, 1997).

**Figure 6:**
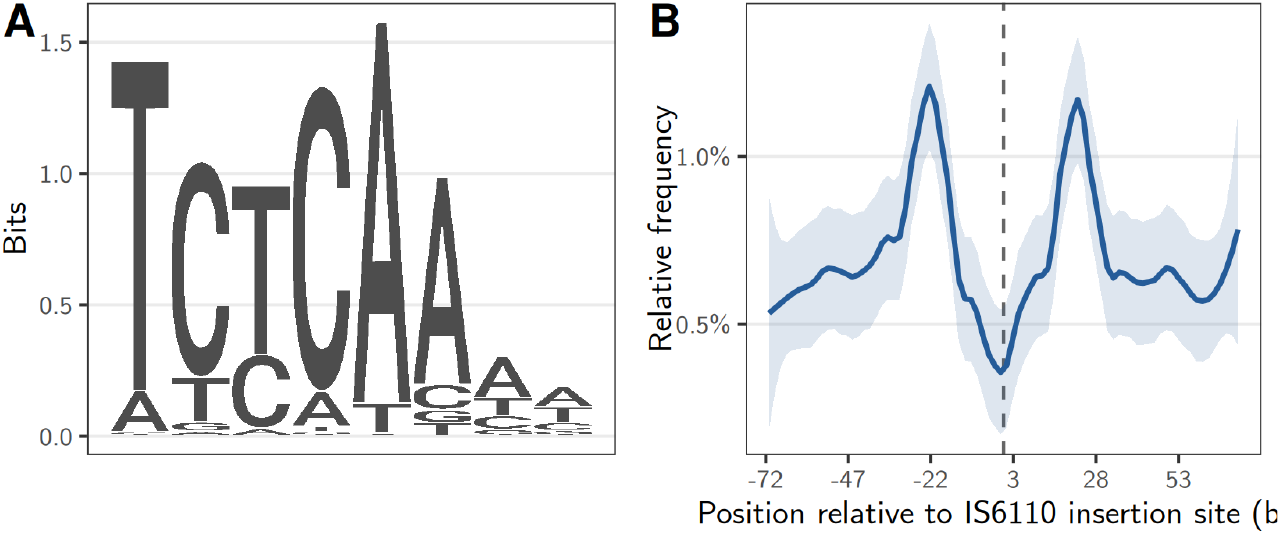
A) Sequence logo for the IS6110 target motif 5’-TCTCAAAW-3’. B) Frequency of motif occurrence relative to the IS6110 insertion site (dashed line).

Given the evidence for target site preferences, we wondered whether an increased density of target motifs contributes to the formation of IS6110 hotspots. To investigate this, the 5’-TCTCAAAW-3’ motif was annotated in the full MTBC0 genome, and its occurrence was counted in all genic and intergenic regions. Of 7361 annotated regions, 1,462 (20%) contained at least one high-confidence match (FIMO *p* < 0.001). Dividing the genome into four categories of IS6110 “hotness”, we observed an increased density of the target motif in “hotter” regions (Fig. S13). Using negative binomial regression to formally test for an enrichment, we inferred median rate ratios ranging from 2.52 to 3.19 over a progression of hotspot thresholds. Effect sizes increase at stricter hotspot thresholds, though with wider credible intervals due to the smaller number of hotspot regions (Fig. S13). These results indicate that hotspot regions contain more IS6110 target motifs per unit length than comparable non-hotspot regions, consistent with a modest but genuine contribution of target motifs to hotspot formation. Hotspots thus tend to be genomic regions that both present adequate target sites and tolerate repeated disruption with no or little fitness cost.

## Conclusion

IS6110 has been studied for more than 30 years, yet much of the literature has taken a pragmatic genotyping perspective or focused on the potential functional consequences of individual insertions. The possibility that high IS6110 copy numbers may benefit the bacterium by increasing genome plasticity and adaptability has been repeatedly proposed (e.g. Gonzalo-Asensio et al., 2018; McEvoy et al., 2007), but evidence for this scenario is scant. As an alternative scenario, we suggest that the distribution of IS6110 across the MTBC and within genomes reflects niche constraints rather than adaptive benefit. A suitable niche for IS6110 is defined by two complementary properties: it provides a favourable target for transposition while imposing little or no fitness cost on the host. Hotspots may emerge where IS6110 is both more likely to insert and more likely to be tolerated once inserted. High copy numbers can then arise through chance insertions into such regions, which both persist for longer and provide additional opportunities for subsequent transposition. Singleton insertions, on the other hand, are consistent with insertions at sites offering no fitness advantage, most of which are eventually purged. This framework does not imply that positive selection of individual insertions never occurs, but rather that adaptive insertions may be side effects rather than the primary reason for the maintenance of IS6110 in the MTBC.

## Methods

### Reference-free detection of insertion sequence polymorphisms

For this study we developed detettore6110, a method to infer insertion sequence polymorphisms from short reads that does not require mapping against a reference genome (Fig. 1). The principle underlying detettore6110 is that an insertion site can be characterized and compared to other insertion sites through the identity of the sequences flanking the insertion rather than its position relative to a reference. In a first step, reads are mapped against an IS consensus sequence, and reads overlapping the beginning or the end of the IS are extracted. The flanking sequences (FS) are then clustered based on a seed sequence immediately adjacent to the insertion breakpoint, requiring an exact match across the first k nucleotides (default 20 bp). Finally, a consensus sequence is called from each FS cluster. For an intact element, this should result in two consensus flanking sequences (cFS), one on the 5’ and one on the 3’ side of the IS.

After flanking sequences have been called in single samples, detettore6110 can combine these results and summarize them in presence-absence matrices where each row is a sample and each column a unique flanking sequence (uFS, Fig. 1B). To decide whether a 5’ cFS from strain A and a 5’ cFS from strain B represent the same uFS, detettore6110 uses the same seed-based strategy as for FS clustering. Specifically, the k-bp sequences immediately adjacent to the insertion breakpoints are compared; if they differ by at most m positions (default 4), the cFSs are assigned to the same uFS. Avoiding full-length sequence alignments substantially improved computational efficiency when comparing thousands of strains.

The output files of detettore6110 contain several quality metrics. For the cFS in a single sample, we included the number of supporting reads, a Z-score indicating whether this number is an outlier relative to the others, and the entropy of the sequence alignment of both the FSs and the reads extending into the element. The entropy of an alignment is defined as the average site entropy, calculated using Shannon’s formula, of the aligned reads. A perfect alignment has an entropy of zero, so this value can be used to check whether the reads in an FS cluster are similar beyond the seed length used.

### Benchmarking detettore6110 with simulated reads

To infer the sensitivity and precision of *detettore6110*, we simulated reads from a set of 30 diverse MTBC assemblies (Harrison et al., 2024) at different coverages (20, 30, 40, 50) and read lengths (100, 150). Insertion sequences in the assemblies were annotated with ISEScan (Xie and Tang, 2017), and the 5’ and 3’ flanking sequences were extracted from the assemblies. Three assemblies were excluded at this step (ASM75654v1, ASM73844v1, ASM73847v1), as they contained untypical fragmented IS copies. Given that these assemblies were inferred from short reads, and no fragmented copies were found in the other assemblies, the fragmented IS are most likely artefacts. After running detettore6110 with the simulated data, the inferred cFS were compared to the true flanking sequences from the assemblies in order to infer true positives and false negatives and to calculate sensitivity and precision.

### Sample selection

Based on the benchmarking results, we selected 10,000 MTBC strains with publicly available short-read sequencing data. Starting point was our inhouse database that contains SNP calls and mapping statistics for more than 100,000 MTBC short-read runs. The alignment and SNP calling workflow has been described previously (Windels et al., 2025). For this study, we selected samples with a read length of at least 100 base pairs; a coverage of at least 30-fold and lower than 300-fold; at least 98% of reads assigned to *Mycobacterium tuberculosis*; at least 80% of SNPs with a major allele frequency > 90% ; and a genome-wide read coverage of at least 95%. For lineages with less than 1000 samples available, we included all samples. The other lineages were downsampled randomly in proportion to their representation in our database, such that a final count of 10,000 was reached. The VCF files for these strains were combined into a single file, retaining only sites where the non-reference allele had a frequency > 0.5.

### Phylogenetic inference

To estimate a phylogenetic tree for the 10,000 strains, we first obtained an alignment of single nucleotide polymorphisms from the combined VCF. As existing tools exceeded memory limits, we developed a Python script that uses memory-efficient numerical numpy arrays to create the sequence alignment (available on https://github.com/cstritt/large_variable_alignment). The script also considers the sequencing depth for each strain at each position in order to distinguish missing data from the null or reference allele. A phylogeny was estimated from the resulting alignment using IQTREE v.3.0.1 with the CMAPLE tree search algorithm (–pathogen, Ly-Trong et al., 2024), an approach optimised for the analysis of large numbers of similar sequences. The general time reversible substitution model was used. Because ascertainment bias correction is not available with the CMAPLE option, we adjusted branch lengths (BLs) in the tree as:

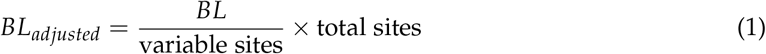

### Ancestral state reconstruction

Ancestral state reconstruction was used in three contexts in this study, when 1) inferring ancestral copy numbers, 2) the numbers of births and deaths on each branch, and 3) the number of independent insertions aggregated per genic and intergenic regions. For each reconstruction we used the maximum parsimony approach implemented in the *asr*_*max*_*parsimony* function of the R castor library (Louca and Doebeli, 2018), which uses Sankoff’s algorithm to infer the smallest number of state changes that leads to the observed distribution (Sankoff, 1975).

For the copy number reconstruction, copy number was estimated for each strain as the lower of the 5’ and 3’ cFS counts. To infer IS6110 births and deaths on each branch, we performed ASR for each of the 10,102 5’ and 10,071 and 3’ uFS, such that for each uFS a presence or absence estimate was obtained for each node of the tree. We then combined these results (still separating 5’ and 3’) by counting the number of 0->1 and 1->0 transitions for each branch of the tree. Finally, we used ASR to identify IS6110 insertion hotspots. For this analysis, we used the 8564 unique insertion sites with mappable reference positions. Because we were interested not in strict parallelism at specific positions but in the parallel disruption of genic and intergenic regions, we aggregated reference positions into a matrix of presence/absence in 1332 regions affected by IS6110. Ancestral states were then inferred for each region separately with maximum parsimony, yielding an estimate for the number of independent insertions into each region.

### Modelling IS6110 birth rates

To model how IS6110 birth rates are affected by variation in copy numbers and branch type (terminal vs. internal), we fitted generalized linear mixed effect models predicting the number of births on a branch as a function of copy number at the parental node (inferred through ASR), branch type, and MTBC lineage. Branch length was incorporated as a log-transformed offset, such that model coefficients are interpreted as effects on birth rates per unit branch length rather than raw counts. Classical phylogenetic comparative methods rely on tip-level traits and incorporate relatedness through a covariance matrix; our approach of modelling branch-level traits instead uses random effects to account for non-independence. Specifically, we included parental node ID as a random intercept to account for branches sharing a parent, and lineage-specific random intercepts and slopes for copy number to allow each lineage to vary in both baseline birth rate and copy-number dependence. We compared models with linear versus log-transformed copy number using AIC to assess the functional form of the copy-number effect. We also tested whether explicitly modelling zero inflation improved fit, which it did not. All models were fitted in R using the glmmTMB package with a negative binomial error distribution to account for overdispersion resulting from the high proportion of branches with zero births.

### Characterization of IS6110 insertion hotspots

A reference genome was required to characterize the genomic distribution of IS6110 and its accumulation in hotspots. We incorporated this option into detettore6110 (Fig. 1A): if a reference genome is provided, detettore6110 uses minimap2 (Li, 2018) to map the 5’ and 3’ cFS against the reference. Then, using the pysam library, detettore6110 extracts pairs of 5’ and 3’ cFS that overlap by a specified number of base pairs that corresponds to the expected size of the target size duplication (3 or 4 bp in the case of IS6110). In this way, insertion sites at base pair precision can be obtained. If an annotation is provided, detettore6110 further outputs the genic or intergenic region into which the IS inserted. As a reference we here used the reconstructed ancestral reference genome MTBC0 (Harrison et al., 2024). To obtain the gene orthologs in the canonical reference H37Rv (ASM19595v2), we identified reciprocal best Blast hits as well as synteny in cases where Blast hits were ambiguous, using Blast v.2.12.0 (Camacho et al., 2009).

The number of independent insertions into each region were inferred through ASR, as described above. To quantify the unevenness of IS6110 insertion distribution across genomic regions, we calculated the Gini coefficient from the distribution of independent insertion events as:

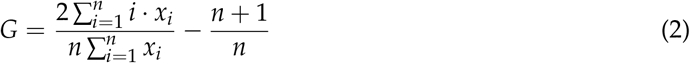

where *x*_*i*_ are the number of independent insertions sorted in ascending order and *n* is the total number of genomic regions. The Gini coefficient ranges from 0 (perfectly uniform insertion distribution) to 1 (complete concentration in a single region).

### Identification of target motifs

To identify potential target site preferences of IS6110, we used the STREME motif finder algorithm v.5.5.9 (Bailey, 2021) to identify motifs enriched in the 8564 unique insertion sites with mappable reference positions. The 5’ and 3’ flanking sequences were joined and the target site duplication was removed such that the insertion breakpoint was located in the middle of the joint sequence. STREME evaluates motif enrichment against a set of background sequences. We generated the same number of background as foreground sequences by randomly extracting regions from the reference, requiring that lengths of sequences are matched. 10 different sets of random background sequences were generated to assess the robustness of the enriched motifs. Then STREME was run with the parameters minw 4, maxw 12, thresh 0.05 and nmotifs 10.

To test whether the top motif is enriched in hotspot regions, the motif was annotated in the MTBCO genome with FIMO v.5.5.9 (Grant et al., 2011), using a relaxed P-value threshold of 0.001 because of the lower scores achieved by a motif with degenerate positions. The number of occurrences was then counted in each region and modelled as function of hotspot status (yes/no) using negative binomial regression, with region length as an offset and region type (genic, intergenic) as a covariate. For each of a progression of hotspot thresholds (10, 20, 40, 60, 80 independent insertion events), 100 independent random samples of non-hotspot regions were drawn and a separate model fitted to each data set, yielding 100 rate ratio estimates per threshold.

## Acknowledgements

We wish to thank Fabrizio Menardo, … for their feedback on the manuscript, as well as the members of the Gagneux group for the many helpful discussions. We also greatly appreciate the work of sciCORE team at the University of Basel, who run their HPC cluster smoothly.

## Data and Code availability

The full workflow of the study from raw data to the figures and tables in the article is provided as a Snakemake workflow on https://github.com/cstritt/MTBC_IS6110-poly. The tool for the reference-free detection of insertion sequence polymorphisms is available on https://github.com/cstritt/detettore6110. The short read data for the 10,000 strains used in the study are publicly available on the NCBI/ENA platforms. A SQlite database with reads mapping to IS6110 insertion breakpoints is available on …; to avoid the trouble of downloading full fastq files, the analysis can be replicated starting from the 4 Gb SQlite file.

## Funding

This work was funded through grants from the European Research Council, grant number 883582, and the Swiss National Science Foundation, grant numbers 10000213, 10001893 and CRSII5_213514.

## Conflict of Interest Disclosure

The authors declare that there are no conflicts of interest.

## Notes

### Competing Interest Statement

The authors have declared no competing interest.

## References

Alonso H, S Samper, C Martín, and I Otal (Dec. 2013). Mapping IS6110 in high-copy number Mycobacterium tuberculosis strains shows specific insertion points in the Beijing genotype. BMC Genomics 14, 422. doi: 10.1186/1471-2164-14-422.

Antoine R, C Gaudin, and RC Hartkoorn (Sept. 3, 2021). Intragenic Distribution of IS 6110 in Clinical Mycobacterium tuberculosis Strains: Bioinformatic Evidence for Gene Disruption Leading to Underdiagnosed Antibiotic Resistance. Microbiology Spectrum 9. Ed. by Lainhart W, e00019–21. doi: 10.1128/Spectrum.00019-21.

Bailey TL (Sept. 29, 2021). STREME: accurate and versatile sequence motif discovery. Bioinformatics 37. Ed. by Birol I, 2834–2840. doi: 10.1093/bioinformatics/btab203.

Camacho C et al. (2009). BLAST+: architecture and applications. BMC bioinformatics 10, 421.

Cerveau N, S Leclercq, E Leroy, D Bouchon, and R Cordaux (Jan. 1, 2011). Short- and Long-term Evolutionary Dynamics of Bacterial Insertion Sequences: Insights from Wolbachia Endosymbionts. Genome Biology and Evolution 3, 1175–1186. doi: 10.1093/gbe/evr096.

Chandler M and J Mahillon (2007). Insertion sequences revisited. Mobile DNA II, 303–366.

Consuegra J et al. (Feb. 12, 2021). Insertion-sequence-mediated mutations both promote and constrain evolvability during a long-term experiment with bacteria. Nature Communications 12, 980. doi: 10.1038/s41467-021-21210-7.

Craig NL (June 1997). Target site selection in transposition. Annual Review of Biochemistry 66, 437–474. doi: 10.1146/annurev.biochem.66.1.437.

Cunningham CW (July 1, 1999). Some Limitations of Ancestral Character-State Reconstruction When Testing Evolutionary Hypotheses. Systematic Biology 48. Ed. by Omland K, 665–674. doi: 10.1080/106351599260238.

Darmon E and DRF Leach (Mar. 2014). Bacterial Genome Instability. Microbiology and Molecular Biology Reviews 78, 1–39. doi: 10.1128/MMBR.00035-13.

DeJesus MA et al. (Mar. 8, 2017). Comprehensive Essentiality Analysis of the Mycobacterium tuberculosis Genome via Saturating Transposon Mutagenesis. mBio 8. Ed. by Stallings CL. In collab. with Manoil C and Lampe D, e02133–16. doi: 10.1128/mBio.02133-16.

Durrant MG, MM Li, BA Siranosian, SB Montgomery, and AS Bhatt (Jan. 2020). A Bioinformatic Analysis of Integrative Mobile Genetic Elements Highlights Their Role in Bacterial Adaptation. Cell Host & Microbe 27, 140–153.e9. doi: 10.1016/j.chom.2019.10.022.

Fang Z and KJ Forbes (Feb. 1997). A Mycobacterium tuberculosis IS6110 preferential locus (ipl) for insertion into the genome. Journal of Clinical Microbiology 35, 479–481. doi: 10.1128/jcm.35.2.479-481.1997.

Ghanekar K et al. (Sept. 1999). Stimulation of transposition of the Mycobacterium tuberculosis insertion sequence IS6110 by exposure to a microaerobic environment. Molecular Microbiology 33, 982–993. doi: 10.1046/j.1365-2958.1999.01539.x.

Goig GA et al. (Mar. 25, 2025). Ecology, global diversity and evolutionary mechanisms in the Mycobacterium tuberculosis complex. Nature Reviews Microbiology. doi: 10.1038/s41579-025-01159-w.

Gonzalo-Asensio J et al. (Aug. 5, 2014). Evolutionary history of tuberculosis shaped by conserved mutations in the PhoPR virulence regulator. Proceedings of the National Academy of Sciences 111, 11491–11496. doi: 10.1073/pnas.1406693111.

Gonzalo-Asensio J et al. (Apr. 12, 2018). New insights into the transposition mechanisms of IS6110 and its dynamic distribution between Mycobacterium tuberculosis Complex lineages. PLOS Genetics 14. Ed. by Buchrieser C, e1007282. doi: 10.1371/journal.pgen.1007282.

Grant CE, TL Bailey, and WS Noble (Apr. 1, 2011). FIMO: scanning for occurrences of a given motif. Bioinformatics 27, 1017–1018. doi: 10.1093/bioinformatics/btr064.

Harrison LB, V Kapur, and MA Behr (Jan. 4, 2024). An imputed ancestral reference genome for the Mycobacterium tuberculosis complex better captures structural genomic diversity for reference-based alignment workflows. Microbial Genomics 10. doi: 10.1099/mgen.0.001165.

Hawkey J et al. (Dec. 2015). ISMapper: identifying transposase insertion sites in bacterial genomes from short read sequence data. BMC Genomics 16, 667. doi: 10.1186/s12864-015-1860-2.

Ho SYW, S Duchêne, M Molak, and B Shapiro (Dec. 2015). Time-dependent estimates of molecular evolutionary rates: evidence and causes. Molecular Ecology 24, 6007–6012. doi: 10.1111/mec.13450.

Hof AEV et al. (June 2016). The industrial melanism mutation in British peppered moths is a transposable element. Nature 534, 102–105. doi: 10.1038/nature17951.

Hugh BT, EM Sim, T Crighton, and V Sintchenko (Mar. 19, 2025). Emergence of Mycobacterium orygis: novel insights into zoonotic reservoirs and genomic epidemiology. Frontiers in Public Health 13, 1568194. doi: 10.3389/fpubh.2025.1568194.

Iranzo J, M. Gómez, FJ López De Saro, and S Manrubia (June 26, 2014). Large-Scale Genomic Analysis Suggests a Neutral Punctuated Dynamics of Transposable Elements in Bacterial Genomes. PLoS Computational Biology 10. Ed. by Bergstrom CT, e1003680. doi: 10.1371/journal.pcbi.1003680.

Kechin A et al. (Feb. 2023). Selection of IS6110 conserved regions for the detection of Mycobacterium tuberculosis using qPCR and LAMP. Archives of Microbiology 205, 71. doi: 10.1007/s00203-023-03410-5.

Kirchberger PC, ML Schmidt, and H Ochman (Sept. 8, 2020). The Ingenuity of Bacterial Genomes. Annual Review of Microbiology 74, 815–834. doi: 10.1146/annurev-micro-020518-115822.

Koonin EV (Dec. 2016). Splendor and misery of adaptation, or the importance of neutral null for understanding evolution. BMC Biology 14, 114. doi: 10.1186/s12915-016-0338-2.

Kurepina N et al. (1998). Characterization of the phylogenetic distribution and chromosomal insertion sites of five IS6110 elements in Mycobacterium tuberculosis: non-random integration in the dnaA–dnaN region. Tubercle and Lung Disease 79, 31–42. doi: 10.1054/tuld.1998.0003.

Li H (Sept. 15, 2018). Minimap2: pairwise alignment for nucleotide sequences. Bioinformatics 34. Ed. by Birol I, 3094–3100. doi: 10.1093/bioinformatics/bty191.

Lok KH et al. (Nov. 2002). Molecular Differentiation of Mycobacterium tuberculosis Strains without IS 6110 Insertions. Emerging Infectious Diseases 8, 1310–1313. doi: 10.3201/eid0811.020291.

Louca S and M Doebeli (2018). Efficient comparative phylogenetics on large trees. Bioinformatics 34, 1053–1055.

McEvoy CR et al. (Sept. 2007). The role of IS6110 in the evolution of Mycobacterium tuberculosis. Tuberculosis 87, 393–404. doi: 10.1016/j.tube.2007.05.010.

Menardo F, S Duchêne, D Brites, and S Gagneux (Sept. 12, 2019). The molecular clock of Mycobacterium tuberculosis. PLOS Pathogens 15. Ed. by Biek R, e1008067. doi: 10.1371/journal.ppat.1008067.

Merker M, TA Kohl, S Niemann, and P Supply (2017). The evolution of strain typing in the Mycobacterium tuberculosis complex. Strain variation in the Mycobacterium tuberculosis complex: its role in biology, epidemiology and control, 43–78.

Refaya AK, U Vetrivel, and K Palaniyandi (Jan. 2024). Genomic Characterization of IS 6110 Insertions in Mycobacterium orygis. Evolutionary Bioinformatics 20, 11769343241240558. doi: 10.1177/11769343241240558.

Rojas VMR et al. (Oct. 9, 2025). Mycobacterium tuberculosis complex Lineage 1: A neglected cause of tuberculosis. PLOS Neglected Tropical Diseases 19. Ed. by Poonawala H, e0013513. doi: 10.1371/journal.pntd.0013513.

Roychowdhury T, S Mandal, and A Bhattacharya (July 28, 2015). Analysis of IS6110 insertion sites provide a glimpse into genome evolution of Mycobacterium tuberculosis. Scientific Reports 5, 12567. doi: 10.1038/srep12567.

Safi H et al. (Apr. 19, 2004). IS6110 functions as a mobile, monocyte-activated promoter in Mycobacterium tuberculosis: M. tuberculosis IS6110 activates downstream genes. Molecular Microbiology 52, 999–1012. doi: 10.1111/j.1365-2958.2004.04037.x.

Sankoff D (1975). Minimal mutation trees of sequences. SIAM Journal on Applied Mathematics 28, 35–42.

Shitikov E et al. (Oct. 2019). The role of IS6110 in micro- and macroevolution of Mycobacterium tuberculosis lineage 2. Molecular Phylogenetics and Evolution 139, 106559. doi: 10.1016/j.ympev.2019.106559.

Siguier P, E Gourbeyre, A Varani, and M Chandler (2015). Everyman’s Guide to Bacterial Insertion Sequences.

Sonnenkalb L et al. (May 2023). Bedaquiline and clofazimine resistance in Mycobacterium tuberculosis: an in-vitro and in-silico data analysis. The Lancet Microbe 4, e358–e368. doi: 10.1016/S2666-5247(23)00002-2.

Soto CY et al. (Jan. 2004). IS 6110 Mediates Increased Transcription of the phoP Virulence Gene in a Multidrug-Resistant Clinical Isolate Responsible for Tuberculosis Outbreaks. Journal of Clinical Microbiology 42, 212–219. doi: 10.1128/JCM.42.1.212-219.2004.

Stritt C and S Gagneux (Sept. 27, 2023). How do monomorphic bacteria evolve? The Mycobacterium tuberculosis complex and the awkward population genetics of extreme clonality. Peer Community Journal 3, e92. doi: 10.24072/pcjournal.322.

Tanaka MM (Aug. 11, 2004). The Control of Copy Number of IS6110 in Mycobacterium tuberculosis. Molecular Biology and Evolution 21, 2195–2201. doi: 10.1093/molbev/msh234.

Thabet S, A Namouchi, and H Mardassi (June 18, 2015). Evolutionary Trends of the Transposase-Encoding Open Reading Frames A and B (orfA and orfB) of the Mycobacterial IS6110 Insertion Sequence. PLOS ONE 10. Ed. by Manganelli R, e0130161. doi: 10.1371/journal.pone.0130161.

Thierry D et al. (Dec. 1990). Characterization of a Mycobacterium tuberculosis insertion sequence, IS6110, and its application in diagnosis. Journal of Clinical Microbiology 28, 2668–2673. doi: 10.1128/jcm.28.12.2668-2673.1990.

Touchon M and EPC Rocha (Apr. 2007). Causes of Insertion Sequences Abundance in Prokaryotic Genomes. Molecular Biology and Evolution 24, 969–981. doi: 10.1093/molbev/msm014.

Ly-Trong N, C Bielow, N De Maio, and BQ Minh (2024). CMAPLE: efficient phylogenetic inference in the pandemic era. Molecular Biology and Evolution 41, msae134.

Vandecraen J, M Chandler, A Aertsen, and R Van Houdt (Nov. 2, 2017). The impact of insertion sequences on bacterial genome plasticity and adaptability. Critical Reviews in Microbiology 43, 709–730. doi: 10.1080/1040841X.2017.1303661.

Wells JN and C Feschotte (Nov. 23, 2020). A Field Guide to Eukaryotic Transposable Elements. Annual Review of Genetics 54, 539–561. doi: 10.1146/annurev-genet-040620-022145.

Werren JH (June 28, 2011). Selfish genetic elements, genetic conflict, and evolutionary innovation. Proceedings of the National Academy of Sciences 108 (Supplement_2), 10863–10870. doi: 10.1073/pnas.1102343108.

WHO (2025). Global tuberculosis report 2025.

Windels EM et al. (June 2025). Onset of infectiousness explains differences in transmissibility across Mycobacterium tuberculosis lineages. Epidemics 51, 100821. doi: 10.1016/j.epidem.2025.100821.

Wu Y, RZ Aandahl, and MM Tanaka (Dec. 2015). Dynamics of bacterial insertion sequences: can transposition bursts help the elements persist? BMC Evolutionary Biology 15, 288. doi: 10.1186/s12862-015-0560-5.

Xie Z and H Tang (Nov. 1, 2017). ISEScan: automated identification of insertion sequence elements in prokaryotic genomes. Bioinformatics 33. Ed. by Hancock J, 3340–3347. doi: 10.1093/bioinformatics/btx433.

Zhang S et al. (2015). Identification of novel mutations associated with clofazimine resistance in Mycobacterium tuberculosis. Journal of Antimicrobial Chemotherapy 70, 2507–2510.

