## Supplementary Figures for "Evolutionary dynamics of the insertion sequence IS6110 in the *Mycobacterium tuberculosis* complex"

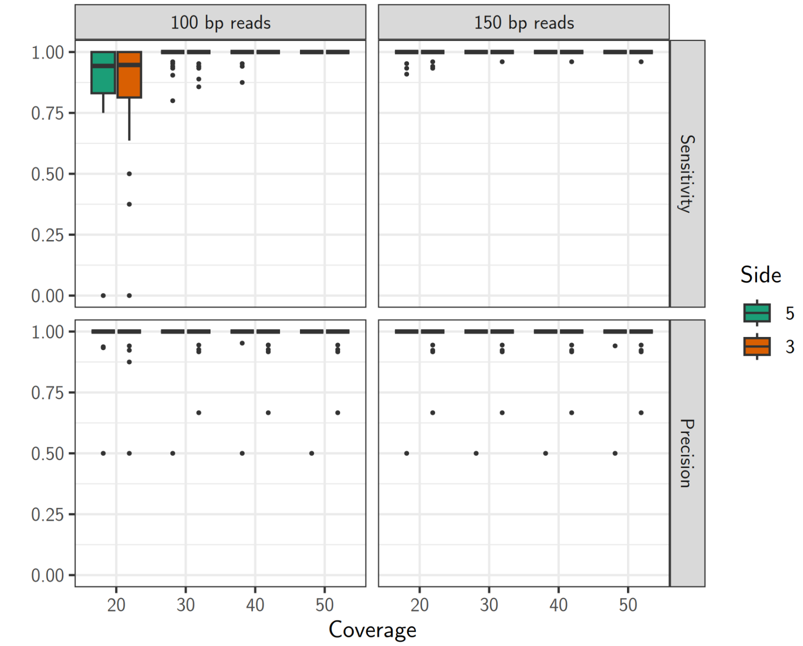

**Figure S1:** *Sensitivity and precision of detettore6110 with simulated reads.*

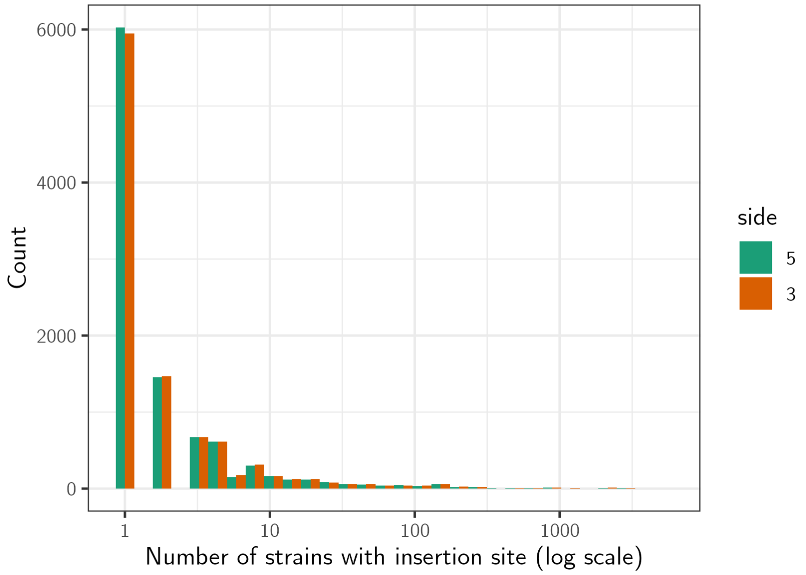

**Figure S2:** *Sensitivity and precision of detettore6110 with simulated reads.*

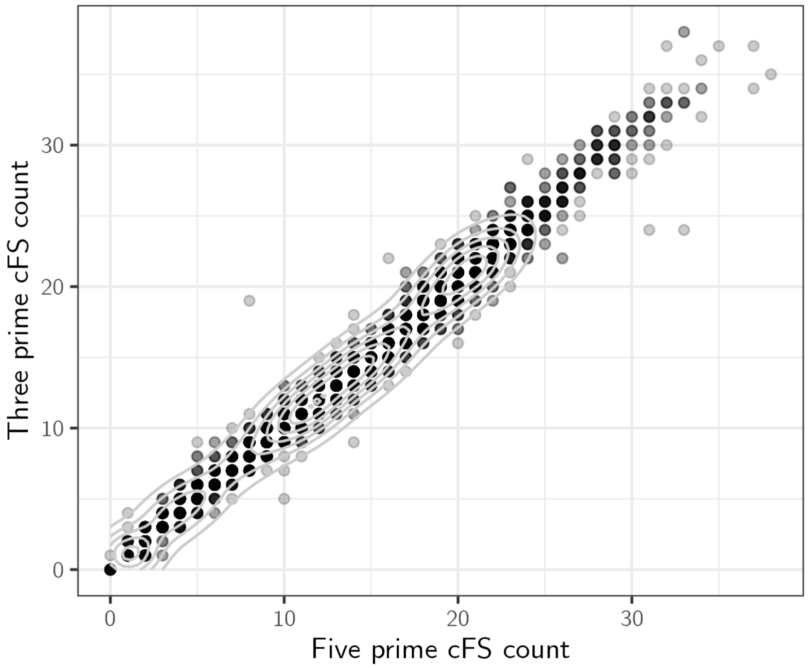

**Figure S3:** *Correlation between IS6110 copy numbers identified from the 5′ and the 3′ consensus flanking sequences (cFS). Each dot is a strain,* $r = 0.995$.

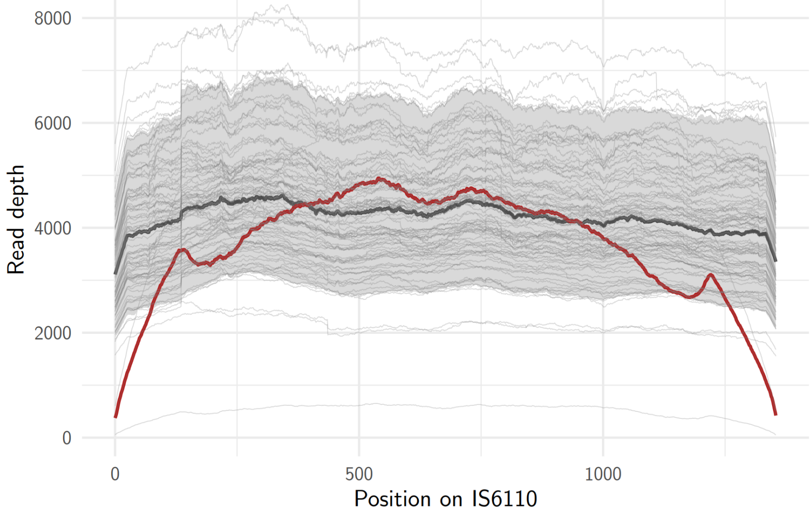

**Figure S4:** *Number of reads mapping to each position of IS6110 for 91 L2 strains from Bioproject PRJNA866200. The strain for which zero copies were inferred (G713449) is highlighted in red. The black line is the median of all strains, the grey ribbon shows the 5-95% interquartile range.*

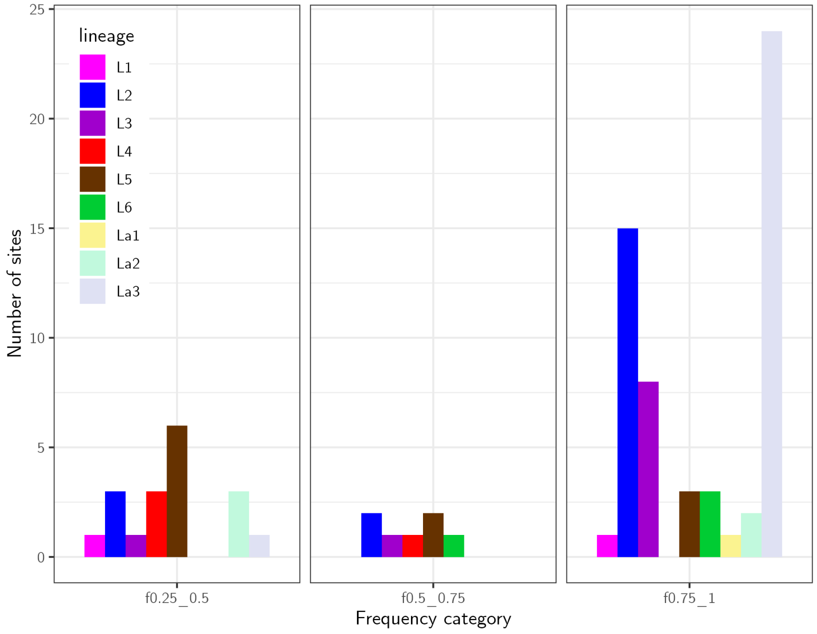

**Figure S5:** *Number of insertion sites that are present at low (0.25-0.5), intermediate (0.5-0.75) and high (0.75-1) frequency in well-represented lineages.*

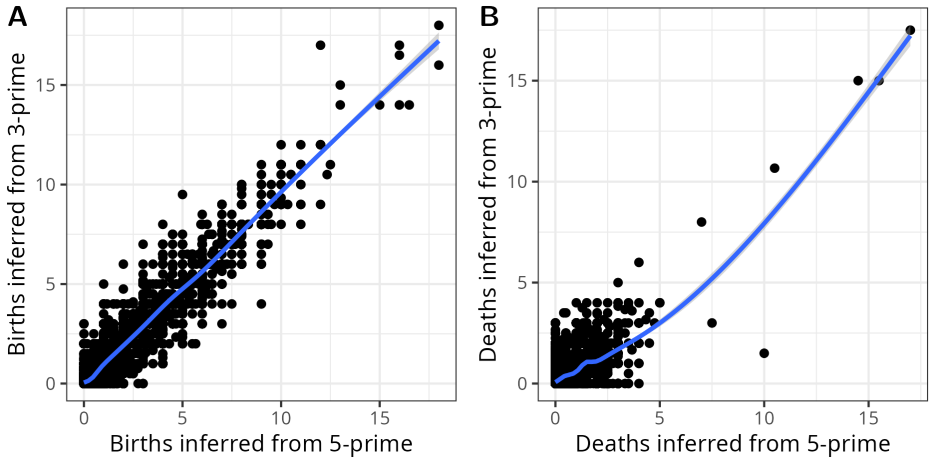

**Figure S6:** *Independent estimates for the number of IS6110 births and deaths on each branch of the tree were obtained from the 5′ and 3′ uFS matrices. The correlation coefficient for the births is* $r = 0.95$*, for deaths* $r = 0.65$*.*

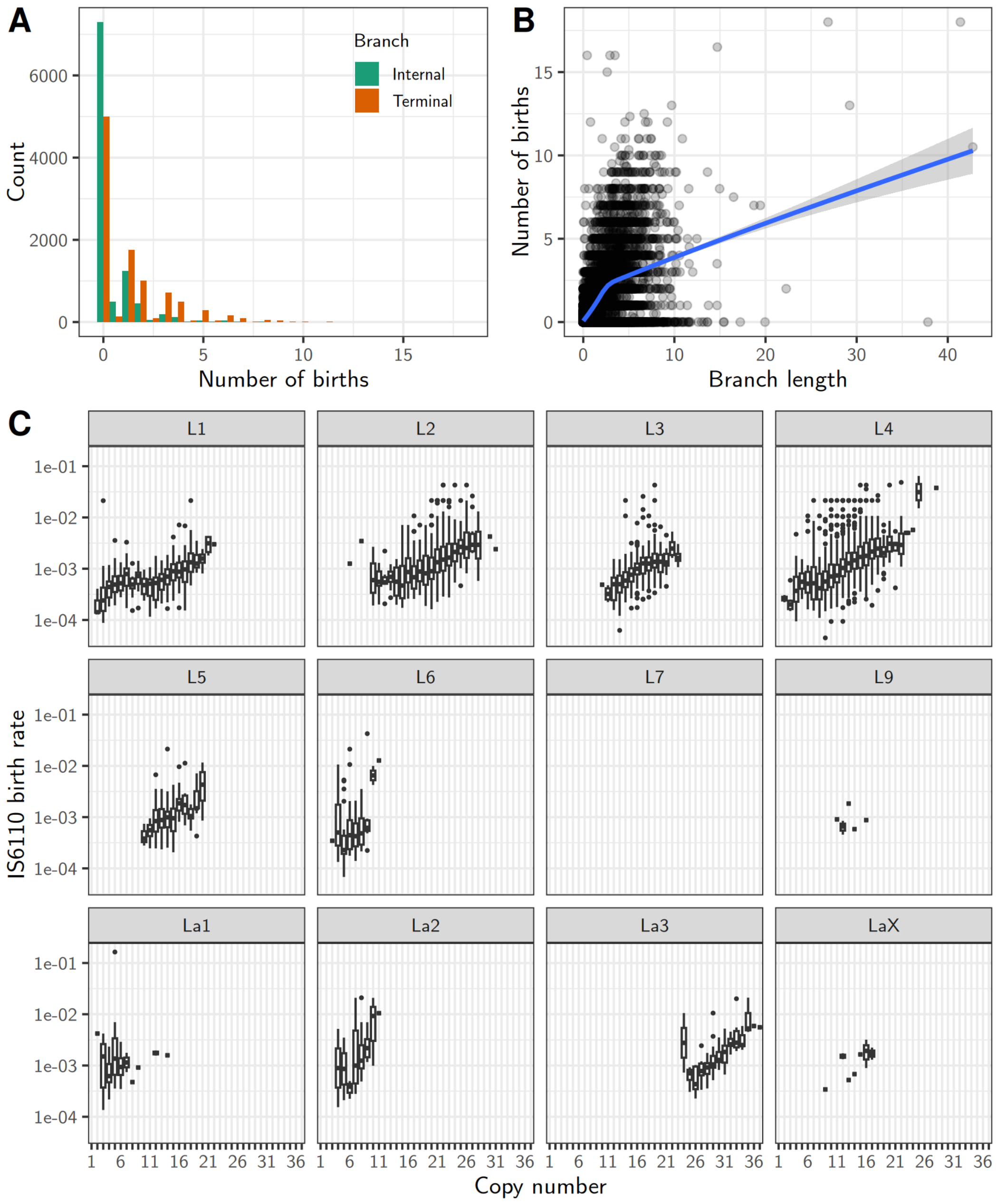

**Figure S7:** *.A) Distribution of the number of births inferred on all branches of the phylogeny, showing that no births were inferred for the majority of branches (12,179). B) Number of births versus branch length, where each dot is one branch. The smoothed curve was fitted with a generalized additive model. C) Distribution of genome-wide IS6110 birth rates. D) Relation between birth rates and IS6110 copy numbers in the different lineages of the MTBC.*

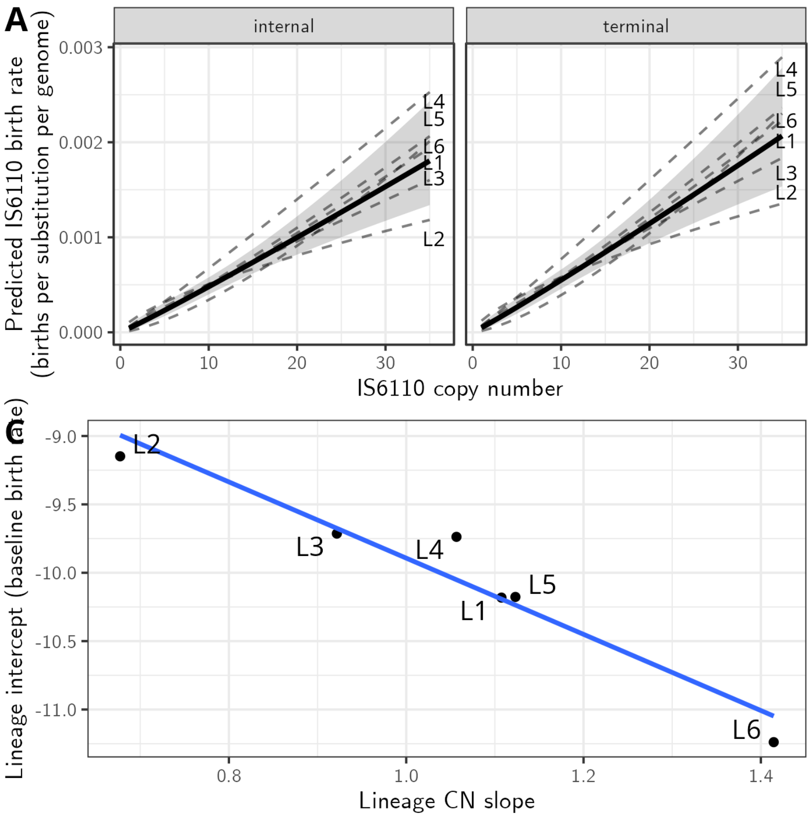

**Figure S8:** *A) Fitted curves with random intercept and slope for each lineage. The population mean effect in bold with shaded confidence interval. B) Negative relationship between random intercepts and slopes ($r = -0.97$).*

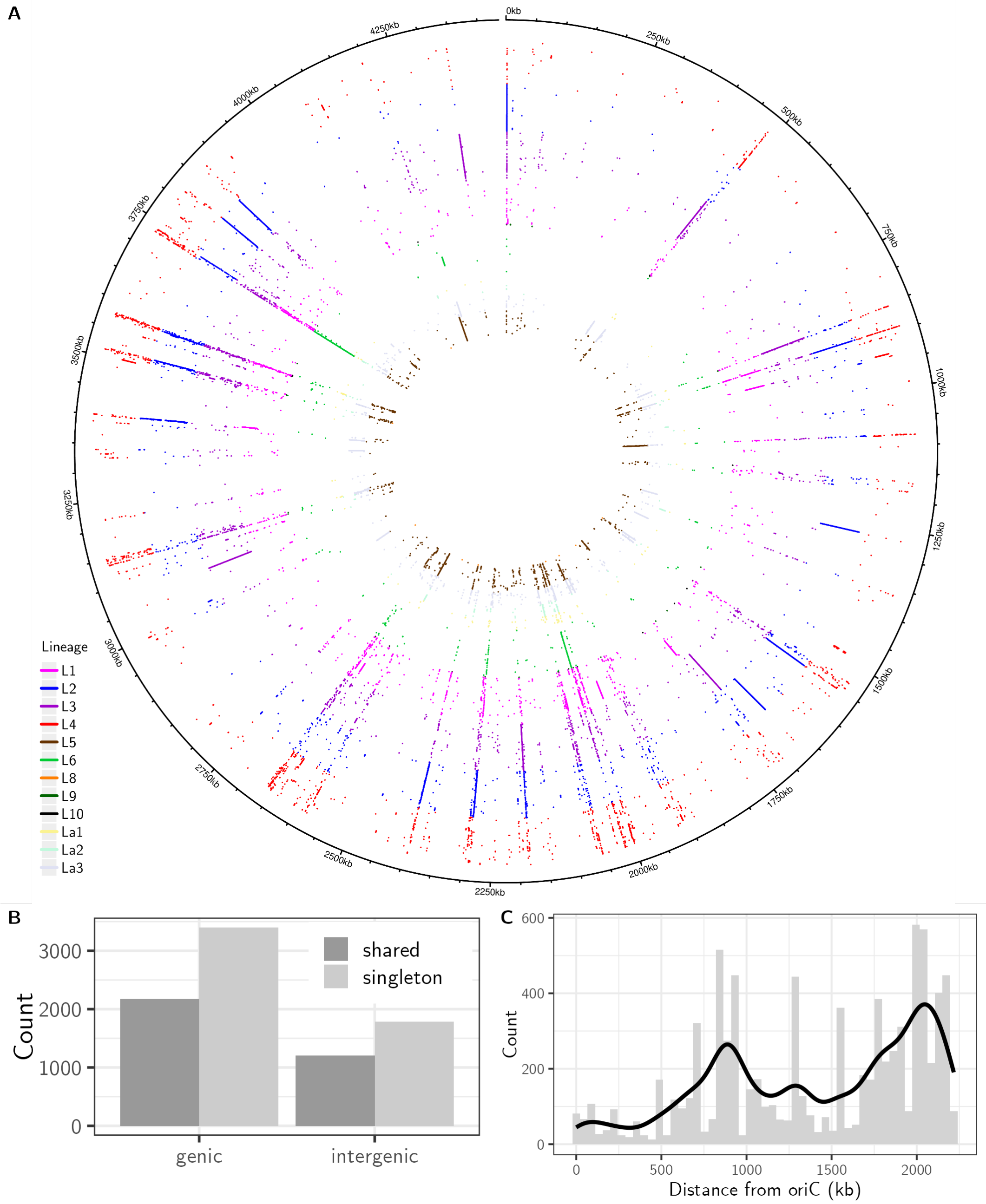

**Figure S9:** *IS6110 insertion sites mapped to the inferred ancestral reference genome MTBC0 (Harrison et al. 2024). Each dot is an IS6110 insertion site, with colors indicating the lineage. To allow visualization, lineages were downsampled to a maximum of 500 strains. The origin of replication (oriC) is indicated by ... 14% of the insertions we inferred could not be mapped to MTB0 and are not represented in the figure.*

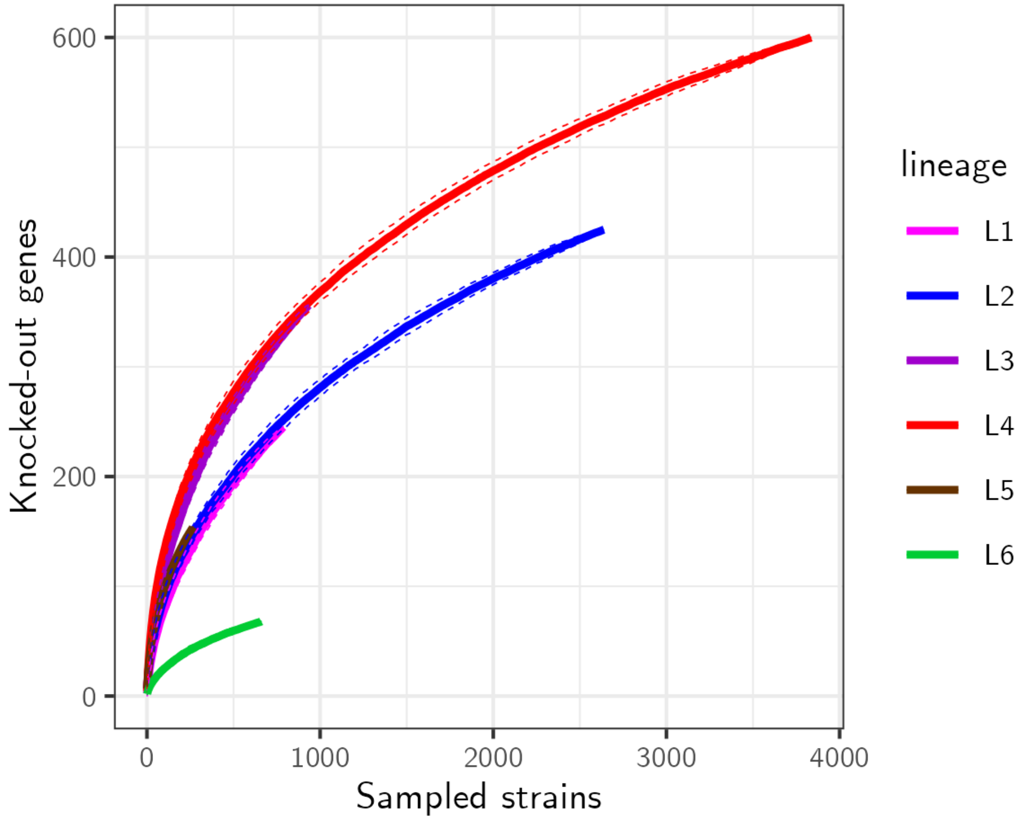

**Figure S10:** *Accumulation curves show the number of unique genes knocked out by IS6110 as additional strains are sampled, separately for the six major MTBC lineages. Solid lines represent the observed accumulation curves, with dashed lines indicating 95% confidence intervals. The continued increase in the number of disrupted genes, including in the well-sampled lineages L1, L2, and L4, indicates that the current sampling has not saturated the diversity of IS6110-disrupted genes.*

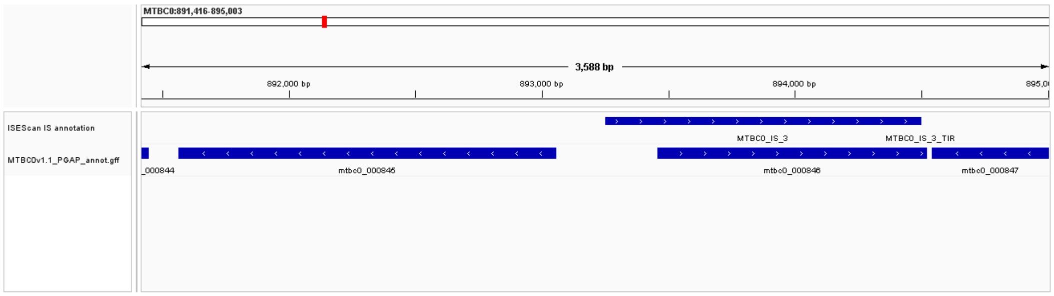

**Figure S11:** *ISEScan annotation of insertion sequences on the upper lane, PGAP gene annotation on the lower.*

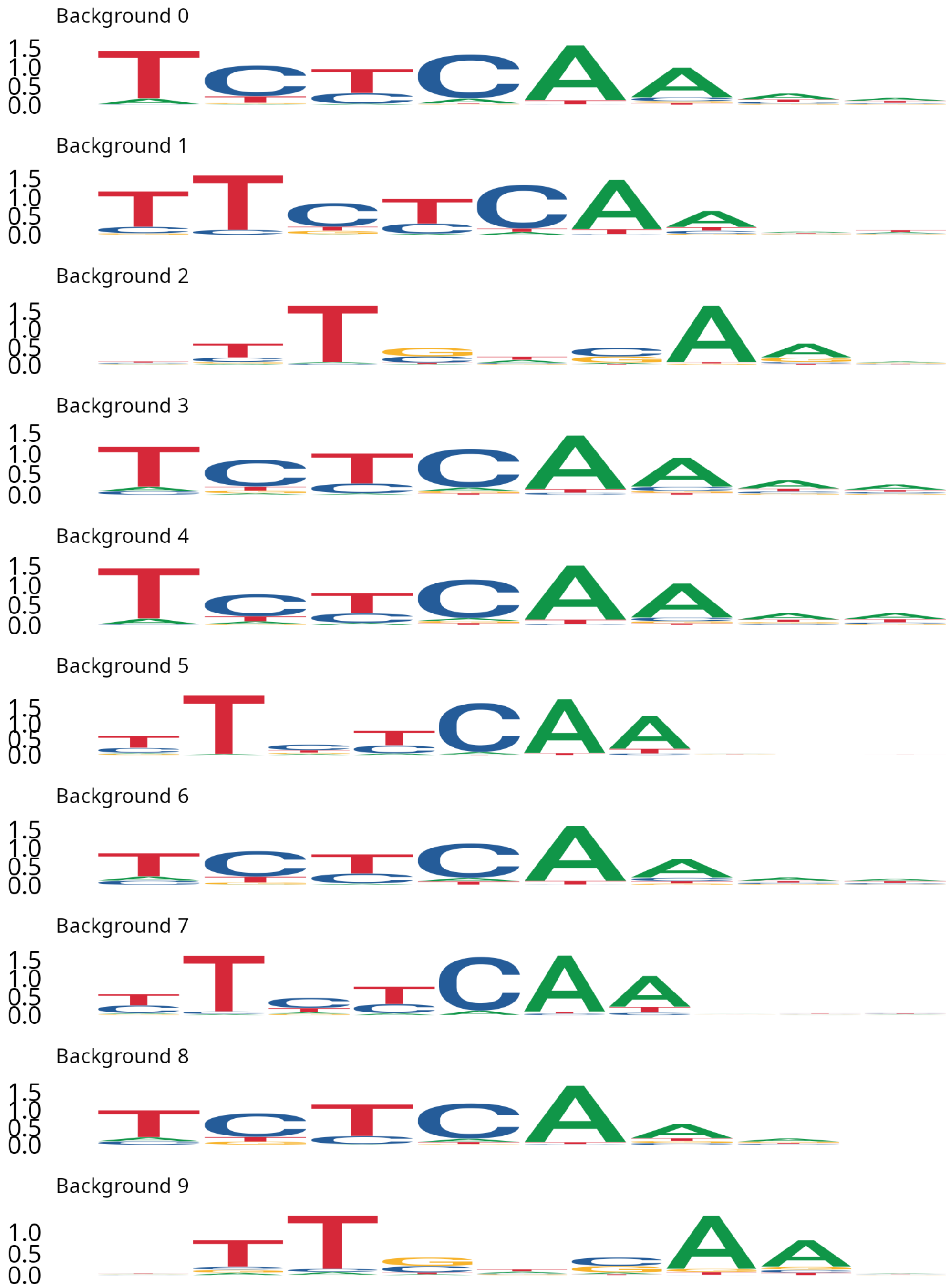

**Figure S12:** *The top target site motifs from ten STREME runs against different random genomic backgrounds. Backgrounds were generated by selecting the same number of random regions as there were foreground sequences, with the same size distribution. As shown in the figure, the top motif was consistently identified, with some variation in degeneracy and motif length.*

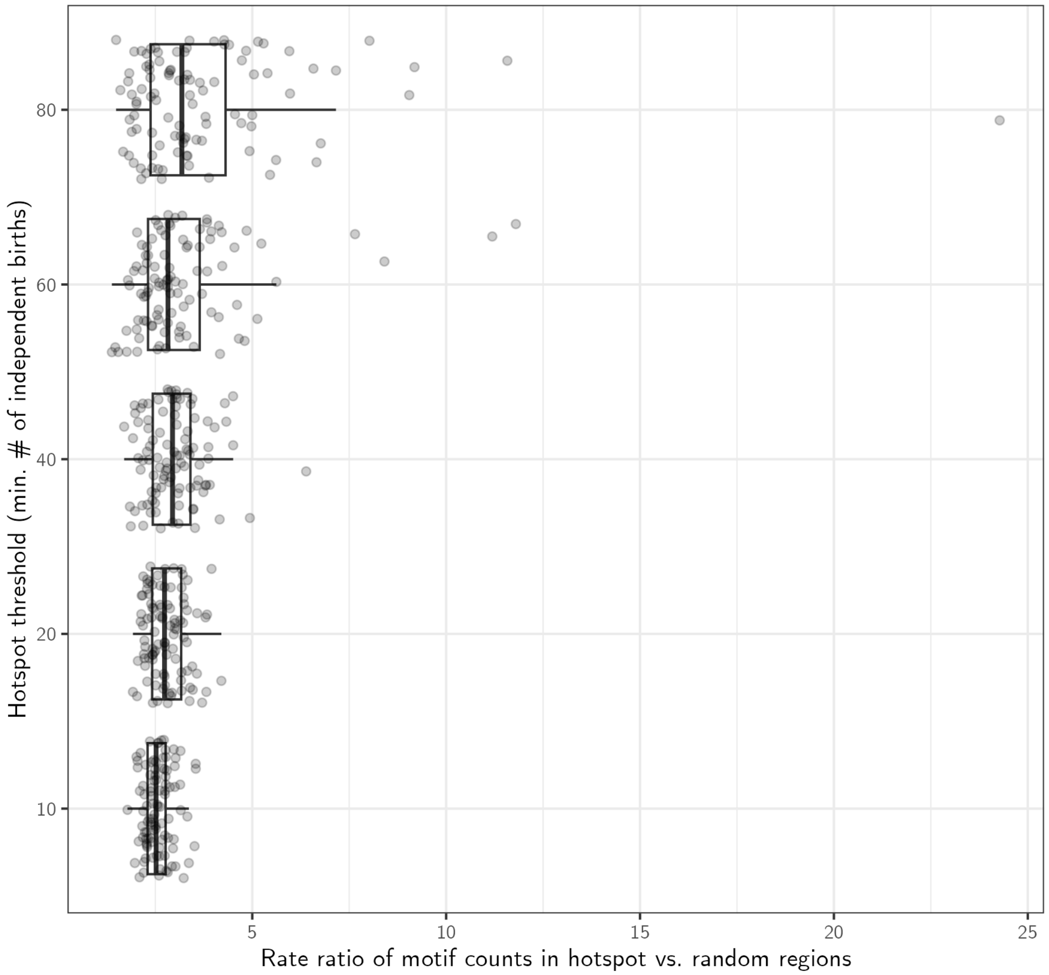

**Figure S13:** *For each of a progression of hotspot thresholds (y-axis), enrichment was tested 100 times with different random regions.*
